# Cryo-EM Structures of *Clostridium perfringens* Enterotoxin Bound to its Human Receptor, Claudin-4

**DOI:** 10.1101/2024.07.11.603128

**Authors:** Sewwandi S. Rathnayake, Satchal K. Erramilli, Anthony A. Kossiakoff, Alex J. Vecchio

**Affiliations:** Department of Biochemistry, University of Nebraska-Lincoln, Lincoln, NE 68588 USA; Department of Biochemistry and Molecular Biology, University of Chicago, Chicago, IL 60637 USA; Department of Structural Biology, University at Buffalo, Buffalo, NY 14203 USA

**Author notes:** Vaxcyte, San Carlos, CA, 94070 USA. Meso Scale Diagnostics, Rockville, MD, 20850 USA.

**Keywords:** Claudins, *Clostridium perfringens* Enterotoxin, Tight Junctions, Cryo-EM, Membrane proteins

## Abstract

Pathogenic strains of *Clostridium perfringens* secrete an enterotoxin (CpE) that causes prevalent, severe, and sometimes deadly gastrointestinal disorders in humans and domesticated animals. CpE binds selectively to membrane protein receptors called claudins on the apical surfaces of small intestinal epithelium. Claudins normally construct tight junctions that regulate epithelial paracellular transport but are hijacked from doing so by CpE and are instead led to form claudin/CpE “small complexes”. Small complexes are building blocks for assembling oligomeric *β*-barrel pores that penetrate the plasma membrane and induce gut cytotoxicity. Here we present structures of CpE in complexes with its native claudin receptor in humans, claudin-4, at 4.0 and 2.8 Å using cryogenic electron microscopy. The structures reveal the overall architecture of the small complex, that the small complex can be kinetically trapped, and resolve its key features; like the residues used in claudin/CpE complex binding, the orientation of CpE relative to the membrane, and CpE-induced structural changes to claudin-4. Further, the structures allude to the biophysical procession from small complex to cytotoxic *β*-barrel pore used by CpE during pathogenesis and the role of trypsin in this process. In full, this work elucidates the structure and mechanism of claudin-bound CpE pore assembly and provides strategies to obstruct its formation to treat CpE-induced gastrointestinal diseases.

## Introduction

*Clostridium perfringens* enterotoxin (CpE) is a 35 kDa two-domain protein toxin secreted by type F strains upon sporulation in the gastrointestinal tracts of humans and domestic animals.^1,2^ CpE causes common foodborne and antibiotic resistant enteritis and enterotoxemia, some of which are severe or fatal in vulnerable populations.^3^ With 1,000,000+ cases of CpE-based food poisoning annually in the United States, it is the third most prevalent with an estimated $400,000,000 economic burden.^4-7^ The large number of cases and high economic burden of CpE is inflated when considering its effects on domesticated animals and livestock.^8^ With no effective vaccine in humans and untreatable enterotoxemia afflicting other animals, therapeutics targeted against CpE could enhance the lives of millions.

The receptors of CpE, claudins, are a family of ∼25 kDa integral membrane proteins that reside on cell surfaces at apical domains of epithelia.^9,10^ Here, claudins self-assemble and connect to the cytoskeleton to form macromolecular ultrastructures called tight junctions that regulate paracellular transport between cell sheets.^11^ In the gut, CpE locates and binds claudins through interactions driven by its C-terminal domain (cCpE; 15 kDa), effectively tethering CpE to cell membranes.^12-14^ Through cCpE, CpE recognizes many claudins but binds well to only a subset of the 27 human subtypes, using a 12 amino acid fingerprint unique to this subset to impart selective and high-affinity binding.^15-17^ Crystal structures of claudins bound to cCpE have shed light on the molecular determinants of these interactions.^15,18-21^ Interestingly, a claudin need not express in the gut to be a CpE receptor, as it has been shown that CpE binds claudins that have low to no expression in these tissues.^22,23^ Homology of claudin structure and sequence in key regions thus plays a role in distinguishing receptors from non-receptors. While bound to claudins, CpE’s N-terminal domain (nCpE; 20 kDa) directs oligomerization and assembly of a transcellular membrane-penetrating *β*-barrel pore with Ca^2+^ selectivity.^24,25^ The first 25 amino acids of nCpE can be removed by trypsin yielding a 32 kDa variant with three-fold higher cytotoxic activity.^26-29^ En masse, cCpE and nCpE coordinate to impart CpE with properties that dissociate tight junctions, breakdown gut barriers, and ultimately damage or destroy epithelial cell sheets.

The claudin-bound CpE *β*-pore that causes epithelial cytotoxicity is predicted to be ∼425 kDa and comprised from six claudin/CpE “small complexes” that associates in 1:1 stoichiometry.^30^ *In vivo* this complex may be short- or long-lived depending on local concentrations of CpE, which are approximated to range from 3-350 nM during infection.^31^ At high concentrations, CpE rapidly kills cells via oncosis while at low concentrations apoptotic signals are activated, each of which induce variable levels of morphological damage to cells and tissues.^3^ We have shown previously that cells lacking other tight junction proteins but expressing a single claudin receptor are susceptible to CpE toxicity and morphological damage, which indicates that structural transitions from small complexes to functional *β*-pores do not require accessory proteins (although in animal tissues other proteins may facilitate larger assemblies).^15,21^ We sought to elucidate the structural basis of the small complex to determine if interactions observed in claudin/cCpE crystal structures were preserved, to reveal the structure and pose of nCpE when CpE is bound to claudins, and to visualize the interfaces available to serve as hubs for *β*-pore assembly. Here we present failed and successful attempts to determine structures of the small complex by X-ray crystallography and cryogenic electron microscopy (cryo-EM), with the latter resulting in the first structures of CpE bound to its endogenous human receptor, claudin-4. Comprehending the interactions, structures, and mechanisms that govern the interplay between claudins and CpE will shed light on this biology; improve current strategies that employ CpE to image, modulate or kill claudin-expressing tissues; and potentially uncover new strategies to inhibit small complex formation to remedy CpE-based gastrointestinal disorders.^32-42^

## Results

### CpE Variant Analyses

Three CpE variants were produced for structural studies that included full-length (CpE), a variant with the first 33 amino acids removed and the DNA level (CpE NΔ33), and trypsinized CpE (CpE_Tryp_), which was prepared by digesting CpE with trypsin.^29^ Although size-exclusion chromatography (SEC) showed all CpE variants to be monomers in solution, blue native-PAGE showed that CpE_Tryp_ migrated differently in-gel compared to CpE and NΔ33 (**Figure 1A**). An X-ray crystal structure of CpE_Tryp_ (PDB ID 8u5f) that was monomeric in solution was found *in crystallo* to form non-crystallographic octamers (**Figure 1A**).^29^ We analyzed CpE_Tryp_ with negative stain EM but large complexes were not observed (**Figure 1A**). Individually, the three variants were mixed with human claudin-4 (hsCLDN-4) and found to form nearly identical 1:1 complexes via SEC and higher molecular weight assemblies by blue native-PAGE (**Figure 1B**). As this showed that removal of CpE’s N-terminus did not affect claudin binding but this had not been quantified before, we used bio-layer interferometry (BLI) to determine the affinity and kinetics of binding. This analysis showed no significant change in CpE variant affinity to hsCLDN-4, with each displaying <10 nM (**Figures 1C** and **S1** and **Table S1**). These hsCLDN-4/CpE variant complexes were used in crystallization trials to determine an X-ray structure of the small complex.

**Figure 1.**
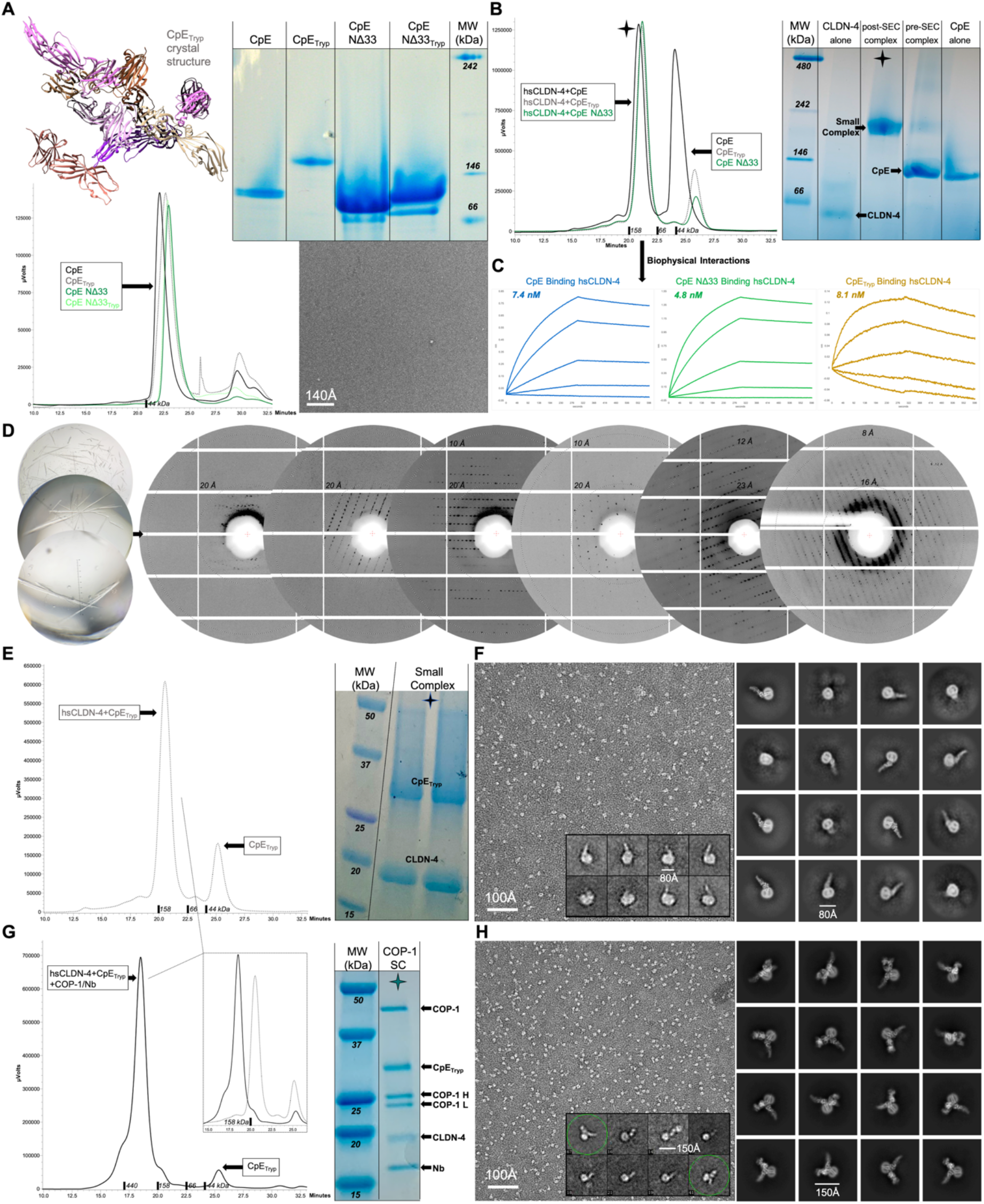
Biochemical, Biophysical, and Initial Structural Analyses of CpE and Small Complexes. (A) CpE variants used in this study as analyzed by SEC, blue native-PAGE, and negative stain EM. The crystal structure of CpE_Tryp_ is shown to illustrate assembly *in crystallo*.^29^ (B) SEC and blue-native PAGE analysis of hsCLDN-4 complexes with three CpE variants. (C) Quantification of biophysical interactions between immobilized CpE variants and hsCLDN-4 using BLI. (D) Crystals of hsCLDN-4/CpE_Tryp_ complexes and progression of X-ray diffraction during attempted optimization. (E) Cryo-EM preparation of hsCLDN-4/CpE_Tryp_ small complexes analyzed by SEC and SDS-PAGE, as well as (F) negative stain EM (left) and cryo-EM (right). (G) Cryo-EM preparation of COP-1 bound hsCLDN-4/CpE_Tryp_ small complexes analyzed by SEC and SDS-PAGE, as well as (H) negative stain EM (left) and cryo-EM (right). In F and H, 2D classes from negative stain appear in the left inset while 2D classes from cryo-EM appear on the right. In G, the inset shows comparative SEC elution profiles from each small complex used for cryo-EM, where COP-1 small complexes elute as larger MW species due to the added 50 kDa COP-1 and 15 kDa Nb.

### X-ray Crystallography of the Small Complex

Using methods that had yielded crystal structures of hsCLDN-4/cCpE (PDB ID 7kp4) we attempted to crystallize hsCLDN-4/CpE.^15^ Crystallization in 96-well format revealed that hsCLDN-4 complexes with NΔ33 and CpE_Tryp_ crystallized more readily than CpE, with polyethylene glycol (PEG) 3350 being the precipitant of choice. Crystals were optimized by altering detergents used to purify hCLDN-4 and incubation temperature. These experiments showed that hsCLDN-4 purified in n-undecyl-β-D-maltopyranoside (UDM) and cholesteryl hemisuccinate Tris salt (CHS) yielded less crystals than in UDM alone and that more crystals grew at 15°C *vs*. lower temperatures. We thus purified hsCLDN-4 in UDM, made complexes with only NΔ33 and CpE_Tryp_, and tested crystallization at 15°C. These experiments showed that complexes with CpE_Tryp_ crystallized in more conditions and grew better appearing and larger crystals (**Figure 1D**). For instance, using the PEG/Ion 96-condition screen we found after one month that 24 *vs*. 12; 24 *vs*. 9; and 15 *vs*. 9 conditions grew crystals of hsCLDN-4 complexes with CpE_Tryp_ *vs*. NΔ33 at 15, 10, and 4°C, respectively. Further crystal optimization strategies included altering the concentrations of protein, salts, PEG3350; the duration of crystallization; as well as scaling to 24-well format to increase crystal size to maximize signal-to-noise in X-ray diffraction. These experiments yielded varied results.

Initial diffraction of crystals scaled to 24 PEG/Ion conditions yielded X-ray diffraction that ranged from >20 to 10 Å resolution (**Figure 1D**). Further optimization through alteration of detergent, protein, salt, and PEG3350 concentration *etc*. yielded only slight improvements to resolution over the next three years. Diffraction was inconsistent from crystal-to-crystal even from those from the same well in the best diffracting conditions, whether they were cryoprotected or not. This indicated that long range disorder in crystals of the small complex was an impeding pathology, possibly due to high solvent content and/or the shapes or dynamics of small complexes in lattices. The best diffraction achieved, ∼7 Å, came from an 11-month-old crystal grown in 19% PEG3350 with 200 mM ammonium phosphate dibasic in space group C2 (cell parameters 517.55, 297.12, 128.71, 90.00, 90.13, 90.00) or P622 (cell parameters 297.02, 297.02, 128.85, 90.00, 90.00, 120.00) (**Figure 1D**). These data could not be indexed or phased well enough to result in a low-resolution map. In sum, despite years of effort, resources, and 1,000+ crystals diffracted from a myriad of conditions, X-ray crystallography had failed to yield a structure of the small complex.

### Cryo-EM of the Small Complex

Concurrently we tried to obtain a structure using cryo-EM. However, because we expected that the ∼55 kDa small complex (23 kDa hsCLDN-4 and 32 kDa CpE_Tryp_) may be too small or dynamic for cryo-EM, we employed two strategies. The first was to use small complexes alone and the second was to add mass and provide a fiducial mark using an anti-hsCLDN-4 synthetic antibody fragment (sFab) to improve cryo-EM amenability.^43^ This sFab, called CpE Obstructing Protein-1 (COP-1), binds hsCLDN-4 and was used recently by us to enable a high resolution structure of the hsCLDN-4/cCpE complex by cryo-EM.^44,45^ After exchanging the solubilizing detergent for hsCLDN-4 from UDM to 2,2-didecylpropane-1,3-bis-β-D-maltopyranoside (LMNG), we made 1 mg of hsCLDN-4/CpE_Tryp_ complexes—henceforth known as small complex(es). CpE_Tryp_ was chosen based on its improved crystallizability and biological role of in activating CpE cytotoxicity.^26^ First, 0.6 mg of small complex was SEC purified resulting in a two-protein complex by SDS-PAGE (**Figure 1E**). Post-SEC, the small complex was imaged using negative stain EM and vitrified and imaged by cryo-EM (**Figure 1F**). To the remaining 0.4 mg of small complex we bound COP-1 and an anti-sFab nanobody (Nb)—henceforth known as COP-1 small complex(es).^46^ This complex was SEC purified resulting in a five-protein complex by SDS-PAGE (**Figure 1G**). Post-SEC, the COP-1 small complex was also imaged using negative stain EM and vitrified and imaged by cryo-EM (**Figure 1H**). For both complexes, negative stain revealed homogenous particles of uniform size that were shaped correctly after 2D classifications; while cryo-EM image analyses revealed 2D class averages with clearly defined secondary structural elements (**Figure 1F** and **1H**). Details of the cryo-EM data collection and processing workflows appear in **Figures S2** and **S3** and final statistics for model building and refinement appear in **Table 1**. These analyses revealed that cryo-EM would successfully yield structures of the small complex.

**Table 1.** Cryo-EM Data Collection, Refinement and Validation Statistics.

|  | hsCLDN-4/ CpE<sub>Tryp</sub> | hsCLDN-4/ CpE<sub>Tryp</sub>/COP-1/Nb |
| --- | --- | --- |
|  | PDB ID: XXXX | PDB ID: XXXX |
|  | EMDB-XXXXXX | EMDB-XXXXXX |
| **Collection & Processing** |  |  |
| Magnification | 129,000x | 105,000x |
| Voltage (kV) | 300 | 300 |
| Electron exposure (e<sup>-</sup>/Å<sup>2</sup>) | 60 | 70 |
| Defocus range (µm) | -1.0 to -2.5 | -0.9 to -2.1 |
| Pixel size (Å) | 0.65 | 1.65 |
| Symmetry imposed | C1 | C1 |
| Number of micrographs | 7,100 | 6,774 |
| Initial particle images (no.) | 2,504,188 | 5,766,325 |
| Final particle images (no.) | 44,683 | 431,680 |
| Map resolution (Å) | 4.00 | 2.83 |
| FSC threshold | 0.143 | 0.143 |
| Map resolution range (Å) | 3.5-46.0 | 2.4-35.8 |
| **Refinement** |  |  |
| Initial models used (PDB ID) | 7kp4 and 8u5f | 8u4v and 8u5f |
| Sharpening B-factor (Å<sup>2</sup>) | -97.9 | -98.4 |
| Model resolution (Å)* | 6.7 / 7.7 | 3.2 / 3.3 |
| FSC threshold | 0.5 | 0.5 |
| Model resolution (Å)# | 4.0 / 4.1 | 2.8 / 2.9 |
| FSC threshold | 0.143 | 0.143 |
| Dimensions (Å) | 65.99, 51.76, 148.81 | 78.57, 139.76, 148.03 |
| Chains | 2 | 7 |
| Atoms | 3,553 | 7,883 |
| Residues | 466 | 1,025 |
| Mean B-factors (Å<sup>2</sup>) |  |  |
| Protein | 142.69 | 138.88 |
| Ligand | N/A | 140.22 |
| Water | N/A | 87.46 |
| R.M.S. deviations |  |  |
| Bond lengths (Å) | 0.004 | 0.003 |
| Bond angles (°) | 0.872 | 0.810 |
| Validation |  |  |
| MolProbity score | 2.32 | 1.75 |
| Clashscore | 27.29 | 8.70 |
| Poor Rotamers (%) | 0.77 | 0.81 |
| Ramachandran plot |  |  |
| Favored (%) | 94.16 | 95.85 |
| Allowed (%) | 5.84 | 4.15 |
| Disallowed (%) | 0.00 | 0.00 |

\* d Fourier Shell Coefficient (FSC) Model 0.50 Masked / Unmasked from Phenix

### Moderate Resolution Cryo-EM Structure of the Small Complex

The final cryo-EM map of the ∼55 kDa small complex was processed from 6,273 quality movies and a total of 44,683 particles was reconstructed to an overall resolution of 4.0 Å per gold-standard Fourier shell correlation (FSC) 0.143 criterion with local resolution estimates from 3.5-46.0 (**Table 1, Figures S2** and **S4**). Although the reported resolution exceeds expectations, the map quality did not resemble a 4 Å map and could not resolve individual side chains but was of sufficient quality to unequivocally model the conformations of both proteins (**Figure 2A**). Density correlating to hsCLDN-4 and CpE were delineated by fitting PDB ID 7kp4 into the map using Chimera then superimposing PDB ID 8u5f onto cCpE and fitting again.^15,29,47^ Each protein was rigid body and real-space refined to yield the final structure (**Figure 2B**).

**Figure 2.**
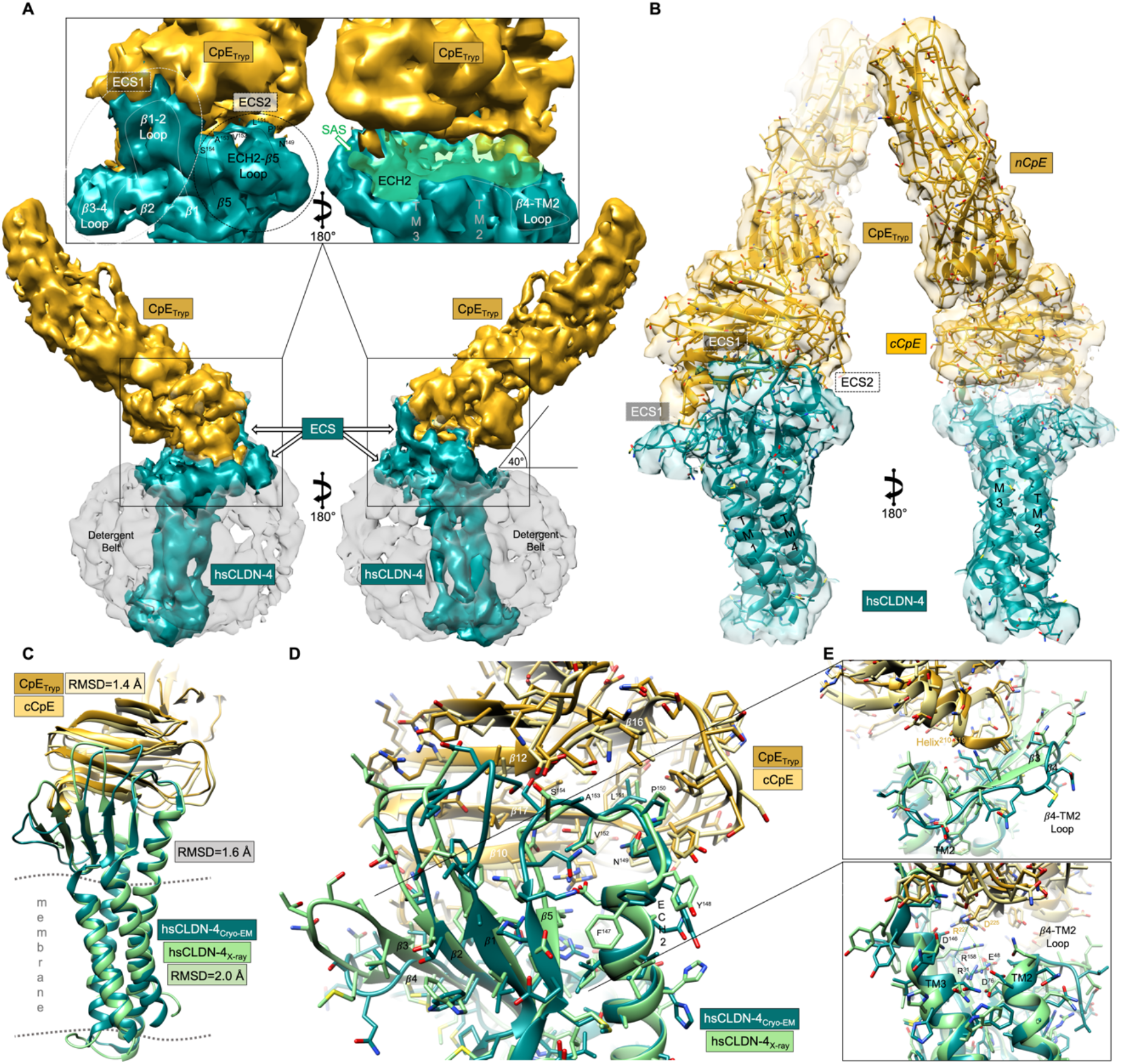
Cryo-EM Structure of the Small Complex. (A) Final 4.0 Å cryo-EM map with hsCLDN-4 (teal), CpE_Tryp_ (gold), and LMNG micelle (grey translucent) densities. Inset shows zoom-in of hsCLDN-4/CpE_Tryp_ interfaces directed by claudin extracellular segments (ECS). Note the solvent-accessible surface (SAS) at the palm region of claudins. (B) The map in A is shown overlaid onto the structural coordinates of the small complex. (C) The hsCLDN-4/CpE_Tryp_ small complex (teal/gold) from cryo-EM superimposed onto the X-ray crystal structure of hsCLDN-4/cCpE (lt. green/lt. yellow) PDB ID 7kp4. (D) Cryo-EM and X-ray structures from C with side chains added show similarities and differences between hsCLDN-4/CpE_Tryp_ and hsCLDN-4/cCpE complexes. Nitrogen, oxygen and sulfur are colored blue, red, and yellow, respectively. (E) Zoom-in of key interfaces and areas of divergence: 1) the *β*3-*β*4 region (top); and 2) *β*4-TM2 loop and triple arginine stack (bottom).

Map and structural analysis showed that CpE_Tryp_ binds to hsCLDN-4 with a distinct pose where extracellular segment (ECS) 2 containing the extracellular helix (ECH) 2-*β*5 loop penetrates a groove between *β*-strands of CpE_Tryp_ in the cCpE domain; ECS1 *β*1-2 loop interacts with the surface of the cCpE domain; and finally, the *β*3-4 loop is displaced by CpE’s 210-219 helix (**Figure 2A** and **2B**). Intermolecular interactions enable formation of a solvent-accessible surface (SAS) at the hsCLDN-4/CpE_Tryp_ interface in the palm region of the hand-shaped claudin structure. These structural features and interfaces influence the CpE binding pose and stabilize it in a single orientation where it relates to the approximated membrane plane at an angle of ∼40°.

Superimposing PDB ID 7kp4, the crystal structure of hsCLDN-4/cCpE, onto the small complex from cryo-EM provides insights into how nCpE affects hsCLDN-4 structure (**Figure 2C**). We calculated an overall root mean square deviation (RMSD) of 1.6 Å between the hsCLDN-4/cCpE structure and the small complex, where comparable chains yielded RMSDs of 2.0 Å and 1.4 Å for hsCLDN-4 and cCpE domains, respectively. This indicated that the structure of hsCLDN-4 was most divergent, and that small complex formation did not greatly alter the conformation of the cCpE domain. Further, this exercise showed that the hsCLDN-4 ECH2-*β*5 loop that contains the NPLVA^153^ motif penetrates the area between *β*16 and *β*17 of cCpE in both structures similarly (**Figure 2D**). However, ECS1 secondary structural elements are changed more significantly. The *β*1-2 loop, the *β*3-4 strands and loop, and the loop connecting *β*4 to transmembrane helix (TM) 2 are increasingly more perturbed the further they are away from ECS2 (**Figure 2E**). In full, despite some structural and conformational similarities between crystal and cryo-EM structures, the larger CpE_Tryp_ alters hsCLDN-4 structure more prominently than the cCpE domain alone in specific regions. The CpE binding pose and structural alterations to claudins observed in the structure of the small complex may affect formation and function of CpE *β*-pores.

### High Resolution Cryo-EM Structure of the COP-1 Small Complex

The final cryo-EM map of the ∼120 kDa COP-1 small complex was processed from 6,627 quality movies and a total of 431,680 particles was reconstructed to an overall resolution of 2.8 Å per gold-standard Fourier shell correlation (FSC) 0.143 criterion with local resolution estimates from 2.4-35.8 Å (**Table 1, Figures S3** and **S4**). Unlike the small complex map, the COP-1 small complex map had high resolution features that resolved secondary structure, side chains, and an LMNG detergent molecule (**Figures 3A** and **S4**). Density for each of the five proteins was defined by fitting PDB ID 8u4v into the map then superimposing PDB ID 8u5f onto cCpE and fitting again.^45^ Each molecule was real-space refined to yield the final structure (**Figure 3B**). The improved resolution of this map, enabled by COP-1s ability to bind and stabilize the small complex during single-particle analysis, allowed us to clearly resolve the small complex with side chain-level precision.

**Figure 3.**
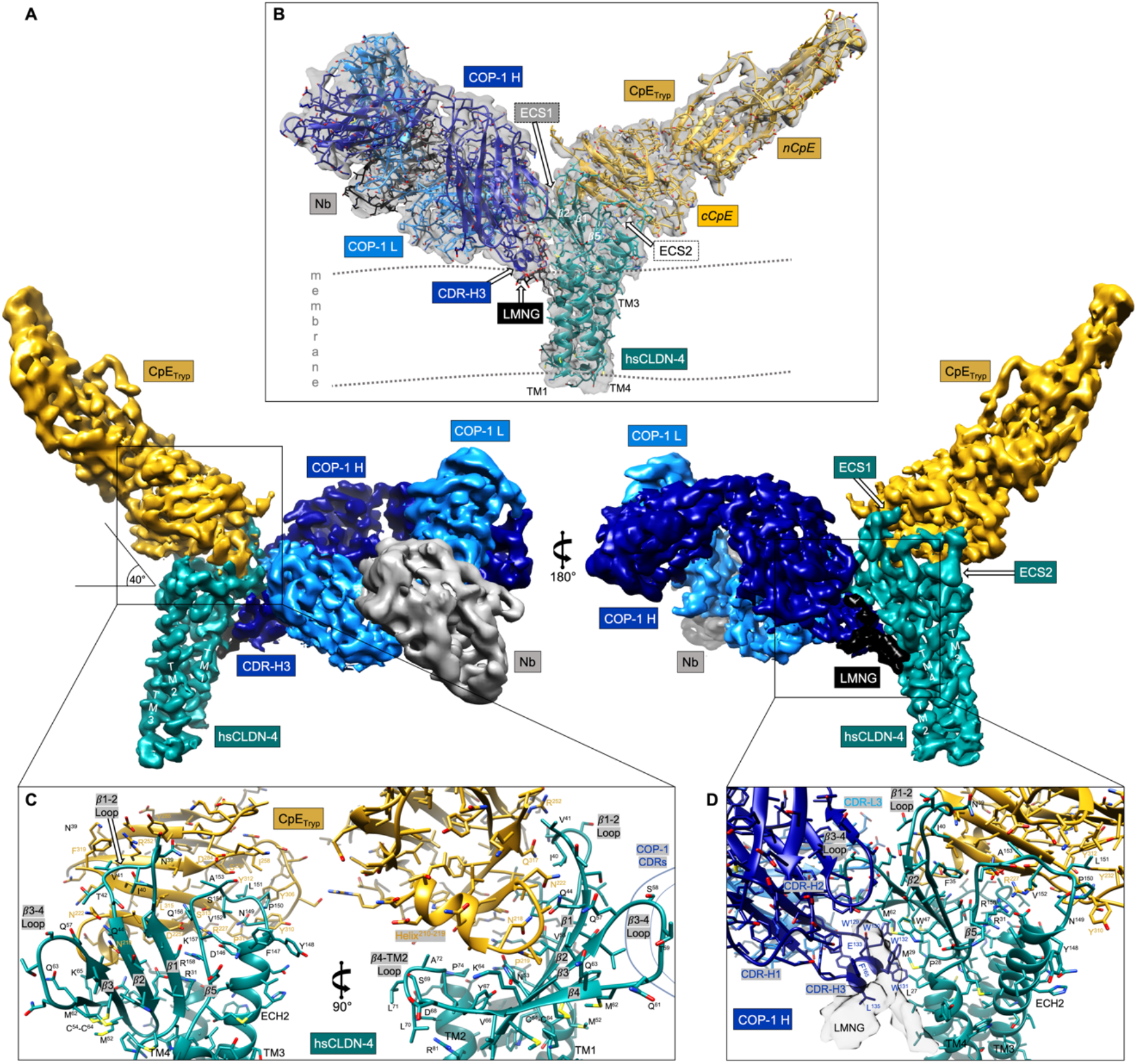
Cryo-EM Structure of the COP-1 Small Complex. (A) Final 2.8 Å cryo-EM map with hsCLDN-4 (teal), CpE_Tryp_ (gold), COP-1 Heavy (dk. blue) and Light Chains (lt. blue), nanobody (grey) and LMNG detergent (black) densities. (B) The cryo-EM map (grey translucent) is shown overlaid onto the structural coordinates of the COP-1 small complex, where protein components are colored as in A. (C) The high-resolution map enables side chain resolution to decipher intermolecular interactions between hsCLDN-4 (teal) and CpE_Tryp_ (gold). Nitrogen, oxygen and sulfur are colored blue, red, and yellow, respectively. (D) The intermolecular interactions between hsCLDN-4 (teal) and COP-1s H (dk. blue) and L (lt. blue) chains are highlighted. The density for an LMNG detergent is shown (white translucent).

Map and structural analysis revealed that the CpE_Tryp_ binding pose was similar to the small complex, where ECS1 and ECS2 interactions with the cCpE domain drive a single stable conformation of CpE (**Figures 3A** and **3B**). Additionally, we found that COP-1 does not interact with any part of CpE_Tryp_, which indicated that natural hsCLDN-4/CpE interactions should be maintained. In whole, we observed three primary regions of hsCLDN-4 used to bind the cCpE domain of CpE_Tryp_ (**Figure 3C**). First, the NPLVA^153^ motif within the ECH2-*β*5 loop penetrates a large pocket created within the *β*16-17 loop of cCpE similar to what we observed in the small complex (**Figure 2D**). Claudin residues from Asp146 to Arg158 form electrostatic, hydrogen, and non-polar interactions with cCpE side chains that span the *β*16-17 loop (**Figure 3C**). Asp146, Asn149, Pro150, Leu151, Ala153, Gln156, and Arg158 all face cCpE to direct intermolecular interactions between hsCLDN-4 ECS2 and CpE_Tryp_. The second region constitutes residues Phe35 to Gln44 within the *β*1-2 loop of hsCLDN-4 (**Figure 3C**). Here, Phe35, Asn39, Val41, and Gln44 form hydrogen bonds and non-polar interactions with *β*10 and *β*17 residues in the cCpE domain. The third region spans *β*3 and *β*4 of hsCLDN-4, which gets displaced by CpE’s 210-219 helix and loop connecting *β*10 (**Figure 3C**). This loop region, focused at Pro219, binds within a groove that is bookended by four claudin side chains, which include Asn53 and Val55 on *β*3 and Gln63 and Lys65 on *β*4. Pro219 penetrates this space between side chain pairs, hovering directly over the 100% conserved Cys54-Cys64 disulfide bond that links *β*3 to *β*4 in claudins—although this disulfide lies on the surface opposite of the cCpE binding interface. The perturbations to hsCLDN-4 structure as a result of CpE_Tryp_ binding observed in the COP-1 small complex coincide with those observed in the small complex. This structure highlights the minimal effect COP-1 has on hsCLDN-4/CpE binding and demonstrates that CpE perturbs claudin structure increasingly from ECS2 through ECS1. Again, CpEs binding mechanism and resulting pose observed here could control the form and function of CpE *β*-pores.

While the cCpE domain of CpE occupies the palm region of the claudin hand, COP-1s heavy (H) and light (L) chains bind to the back of the claudin hand (**Figure 3B**). Using three complementarity-determining regions (CDRs) per chain, COP-1 sandwiches hsCLDN-4s ECS1 between itself and CpE_Tryp_. The *β*3-4 loop of hsCLDN-4 specifically is bound between the H and L chains of COP-1 and is further pressed by CpE’s 210-219 helix (**Figure 3D**). This grip on hsCLDN-4 by COP-1s CDR-H2 and -L3 enables CDR-H3 to do something unique. CDR-H3 is distinctly structured, forming an amphipathic helix that partially penetrates the detergent belt to bind the TM1-*β*1 area of hsCLDN-4 (**Figures 3B** and **3D**). The hydrophobicity of CDR-H3 is confirmed by visualizing the LMNG belt in the map and the presence of an LMNG molecule at the CDR-H3/hsCLDN-4 interface (**Figures 3D** and **S3**). In full, we observed that the interactions driving COP-1 recognition of hsCLDN-4 when bound to CpE_Tryp_ were unaltered compared to those when hsCLDN-4 is bound to cCpE, which we determined previously.^45^ Because COP-1 had minimal effect on hsCLDN-4/CpE interactions, we could accurately compare our two small complex structures to determine CpEs impact to claudin structure.

### Structural Comparison of Small Complexes

Because the hsCLDN-4/CpE_Tryp_ portion of the COP-1 small complex resembled that from the small complex, we compared and analyzed them in greater detail to synthesize our findings. First, we superimposed the hsCLDN-4/CpE_Tryp_ portion of both small complex structures (**Figure 4A**). We found that the overall conformation and pose of CpE_Tryp_ when bound to hsCLDN-4 was similar, yielding an RMSD of 1.5 Å. This was closer than the calculated RMSDs of 1.9 Å overall and 2.0 Å between hsCLDN-4 in the two structures—highlighting again that hsCLDN-4 represented the largest conformational divergence. More detailed analysis of the hsCLDN-4/CpE interface explained this larger divergence in hsCLDN-4. Structural comparison shows that the *β*3-4 loop and *β*4-TM2 loop vary significantly (**Figures 4B**). Because *β*4 links both loops these differences appeared coordinated. As previously shown, the H and L chains of COP-1 grip the *β*3-4 loop to lock it in this conformation, which is distinct and does not occur when not bound to COP-1 (**Figure 3D**). We also found that the *β*1-2 loop was altered between structures but that the ECH2-*β*5 loop was not (**Figure 4C**). The CDRH-2 of COP-1 in binding hsCLDN-4 interacts with both *β*3-4 and *β*1-2 loop side chains but not any on ECS2. These analyses revealed the effect of COP-1 binding to hsCLDN-4 on its structure when in complex with CpE, which adequately explained the observed differences between small complex structures. Overall, despite these changes imposed by COP-1, the overall pose and intermolecular interactions hsCLDN-4 uses to bind CpE were maintained, allowing us to use these structures to predict the biophysical procession from small complex to cytotoxic *β*-pore.

**Figure 4.**
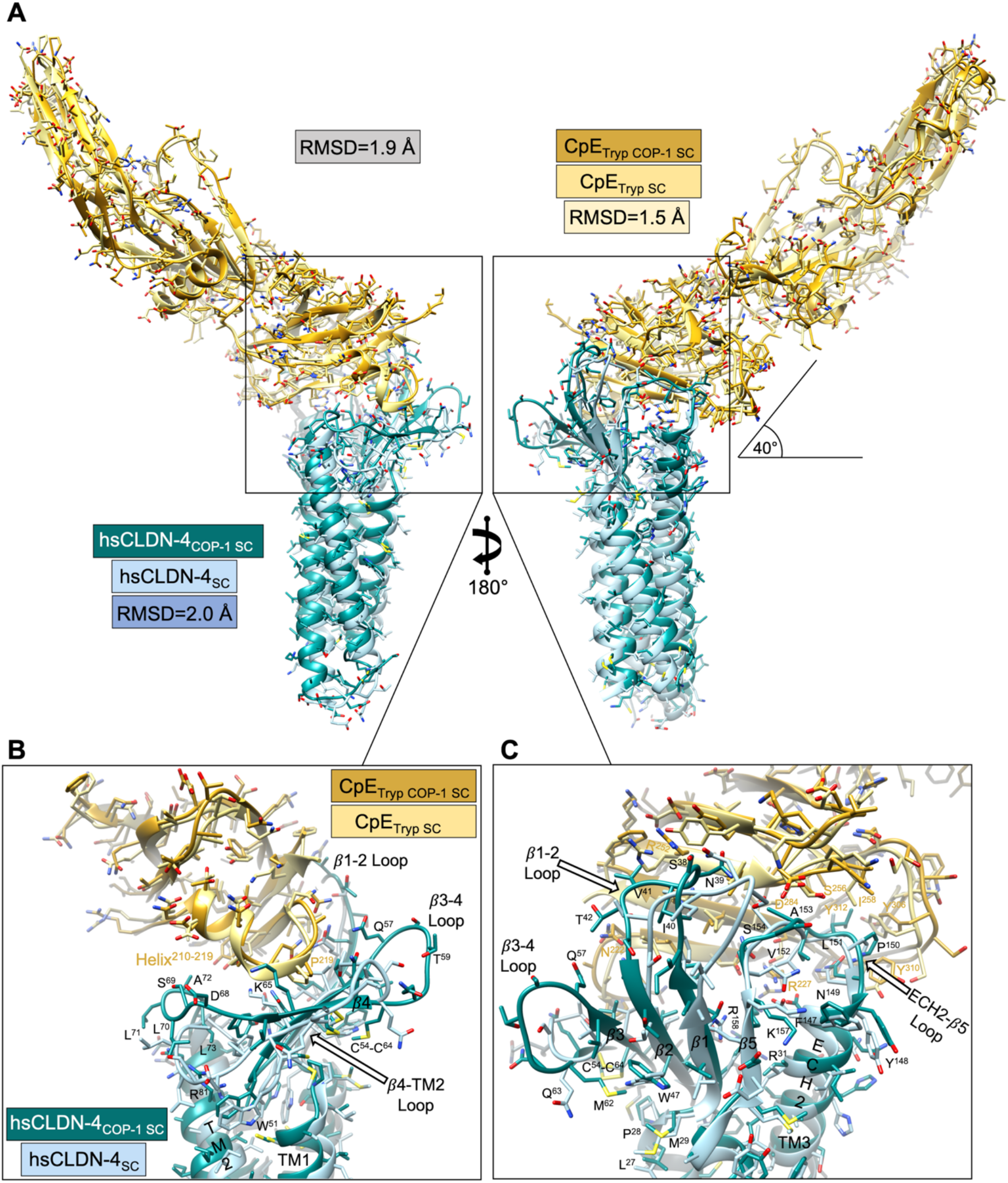
Structural Comparison of Small Complexes. (A) Superimposition of hsCLDN-4/CpE_Tryp_ complexes from the cryo-EM small complex structure (lt. blue/yellow) and COP-1 small complex structure (teal/gold) with RMSD calculations between equivalent chains. (B) Large structural differences arise in the β4-TM2 loop region of ECS1. (C) Progressively larger differences in the conformation of hsCLDN-4 ECS occurs from ECS2 to ECS1, indicating ECS2s role in stabilizing CpE binding and ECS1s resulting disruption due to CpE binding. Note that the observed β3-4 loop (teal) conformation in the COP-1 small complex is influenced by COP-1 binding (which is removed for clarity).

### Small Complex Integration into Functional CpE β-pores

We modeled how our small complex structures might integrate into a functional *β*-pore using computational predictions. Using the sequences of hsCLDN-4 and CpE_Tryp_ we input four to ten copies of each protein into AlphaFold3 (AF3).^48^ We found that tetramers, pentamers, and decamers did not result in physiologically relevant pore-like structures but found that hexamers, heptamers, octamers, and nonamers did (**Figure S5**). For these oligomers, AF3 predicted circular arrangements of small complexes with central pores that extend from the extracellular space where nCpE resides through the membrane and into the intracellular space where claudin four-helical bundles meet (**Figure 5A**). AF3 did not predict a *β*-pore however, but rather what may be the predicted “pre-pore” complex—the oligomeric state of small complexes that assembles on cell surfaces prior to structural reorganization of nCpE to form the membrane-penetrating pore.^49^ These assemblies was deemed relevant due to the hypothesized *β*-pore-forming helix being positioned toward the central pore and Asp48 being positioned toward a CpE/CpE interface.^49,50^ Because the overall orientation of these four to nine small complexes were similar, we focused further investigation on the hexamer because it is the predicted functional unit of cytotoxic CpE *β*-pores.^30^ Superimposing our small complex and COP-1 small complex structures onto a single small complex subunit within the hexameric pre-pore yielded RMSDs of 2.7 Å and 1.4 Å, respectively (**Figures 5B** and **5C**). This indicated that our experimental small complexes could integrate with slight reorganization into oligomeric CpE pre-pores.

**Figure 5.**
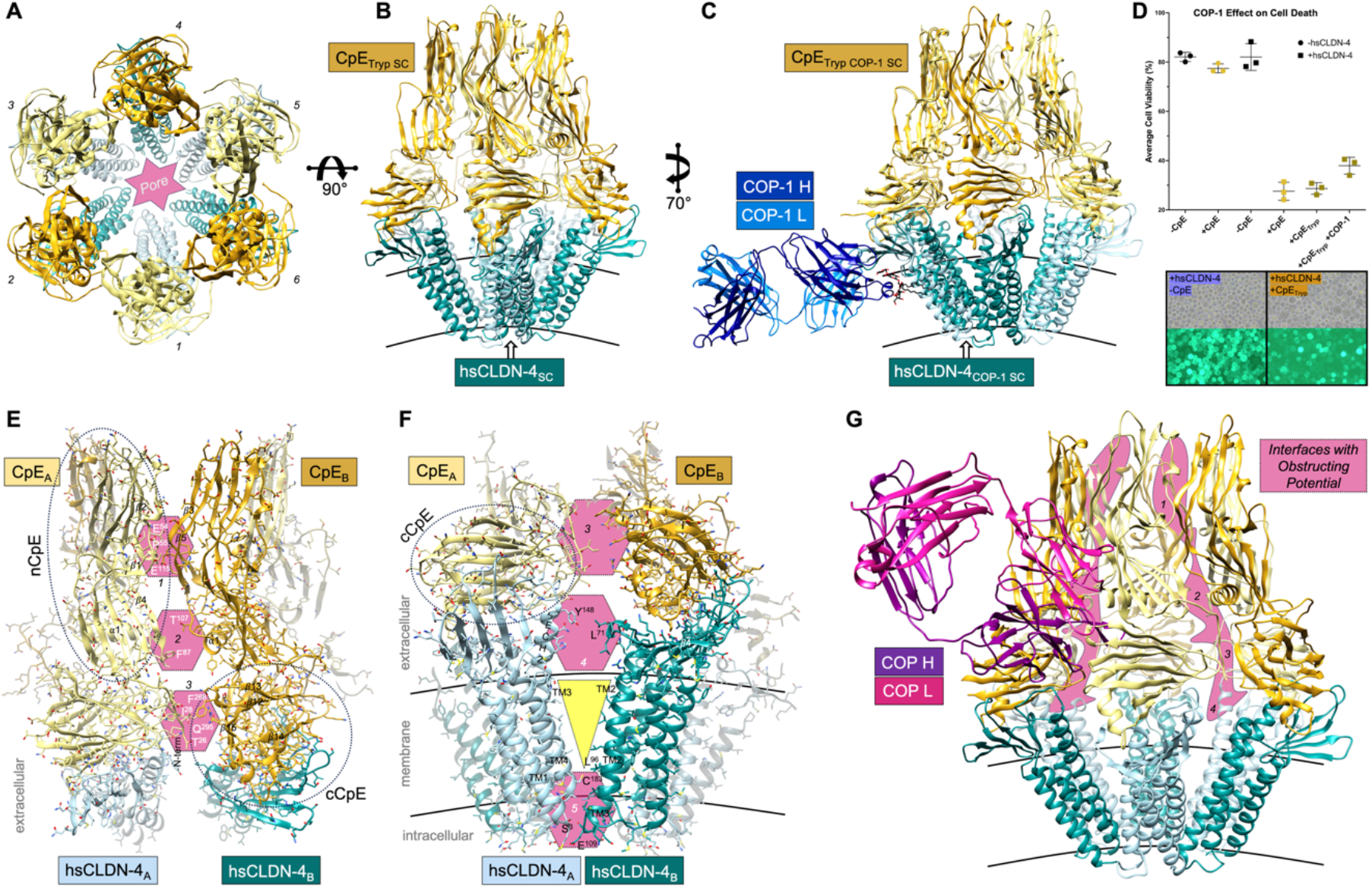
Small Complex Integration, Assembly, and Inhibition of the CpE Pre-Pore. (A) AlphaFold3 prediction of CpE pre-pore using hsCLDN-4 and CpE_Tryp_ sequences as inputs. Each individual hsCLDN-4/CpE small complex is colored teal/gold or lt. blue/yellow. Integration of our (B) small complex (teal/gold) and (C) COP-1 small complex (teal/gold) into a single small complex (lt. blue/yellow) from the AF3 predicted hexameric CpE pre-pore. Note that only minor alterations are needed to fit the experimental small complex structures into the predicted pre-pore complex. (D) CpE-induced cytotoxicity assay. Scatter plot depicts individual cell viability counts from triplicate measurements, with average viability and standard error of the mean, of normal insect cells (circles) or insect cells expressing hsCLDN-4_eGFP_ (squares), untreated (black) or treated with CpE (yellow/gold). Morphological damage and fluorescence changes to cells expressing hsCLDN-4_eGFP_ without (left) or with CpE (right) added is also shown. The interaction interfaces (pink hexagons) of (E) CpE and (F) hsCLDN-4 between two small complexes (teal/gold and lt. blue/yellow), which form the basis of hexameric CpE pre-pore assemblies. (G) Using the AF3 generated CpE pre-pore hexamer, we predict the four interfaces (pink trace) on CpE and hsCLDN-4 that have potential to be targeted and bound to by a hypothetical COP sFab (purple/pink). A COP that binds one or several of these interfaces could potentially obstruct pre-pore assembly and therapeutically prevent CpE β-pore-induced cytotoxicity.

### COP-1 Obstruction of CpE Cytotoxicity

One goal of developing COPs was to discover one capable of binding either claudins or CpE to preclude CpE *β*-pore assembly and thus cytotoxicity. The COP-1 small complex modeled onto the AF3 generated hexameric pre-pore predicted that COP-1 would not bind between any small complex/small complex interfaces (**Figure 5C**). We thus surmised that a COP-1 bound small complex could assemble an oligomeric pre-pore and potentially a cytotoxic *β*-pore normally, so COP-1 would not obstruct CpE cytotoxicity. We tested this hypothesis using a cell based assay where CpE was added to GFP-tagged hsCLDN-4 expressing insect cells and cytotoxicity was monitored using microscopy and quantified with cell staining (**Figures 5D** and **S6**).^15^ We found that insect cells left alone or treated with CpE, and those expressing hsCLDN-4 but not treated with CpE, had average cell viabilities in the range of 77.4-82.1% (mean 80.5%). The 19.5% viability loss for these control wells was attributed to normal time- or baculovirus-induced cell death. We then found in hsCLDN-4 expressing cells that adding CpE or CpE_Tryp_ resulted in average cell viabilities of 27.6 and 28.7%, respectively, indicating that CpE decreased viability by 52.9 and 51.9% more than controls. Lastly, after adding two moles excess of COP-1 just before adding CpE_Tryp_, we measured average cell viability to be 37.9%. This result was a 42.6% decrease compared to controls and 10.4 and 9.4% higher viability compared to CpE and CpE_Tryp_ treated cells (**Figure 5D**). Analysis of the morphological damage to insect cells in the presence of COP-1 showed that there was an absence of very large cells and generally more regularly shaped cells (**Figure S6**). These results confirmed aspects of the CpE pre-pore modeling. But because no therapeutic exists to treat food-borne CpE gastrointestinal disease in humans, we investigated our models of CpE pre-pore assembly in greater detail.

### CpE Pre-pore Assembly

Structural analysis of the AF3 generated hsCLDN-4-bound CpE pre-pore revealed five potential Interaction interfaces that may facilitate pre-pore assembly. Specifically, the models show three sets of interaction interfaces exist between adjacent CpEs in extracellular space (**Figure 5E**). At nCpE, the first constitutes interactions between the *β*1-2 linker region from one subunit and *β*3 and *β*5 from another, while the second comprises the *β*4-α1 linker region from one unit interacting with the α1-*β*5 linker from another. The third interface constitutes interactions between CpE’s N-terminus and cCpE’s *β*12-13 loop region and *β*15. Because we input the sequence of CpE_Tryp_ into AF3 with its 26 amino acid shorter N-terminus, it was interesting to find a potential interaction between this area and other parts of CpE.

The models also show two interaction interfaces that occur between adjacent claudins that may facilitate CpE pre-pore assembly (**Figure 5F**). The first occurs in extracellular space between the ECS2s ECH2 and the β4-TM2 loop of ECS1. The second occurs intracellularly and constitutes a joint TM1-TM4 interface associating with a joint TM2-TM3 interface on a neighboring protomer. The adjacent claudins are bound most closely at the intracellular domain then splay outward moving to extracellular space forming a “V” shape. These interactions require adjacent claudins to tilt at 30° within the membrane creating conical shaped claudin dimers. The conical shape of claudins creates six dimeric interfaces that could influence the shape and fluidity of the plasma membrane prior to CpE *β*-pore formation. In sum, the small complex interaction interfaces create targetable ares for sFabs to bind and obstruct pre-pore formation, these molecules may hold potential as therapeutics (**Figure 5G**).

## Discussion

Although the receptors of CpE were discovered 25+ years ago and subsequent studies shed light on the interactions and functional consequences of CpE binding to claudins, the structural basis of CpE-based cytotoxicity had to be inferred from structures of claudins bound to cCpE determined in the last 10 years. This study provides the first structures of the small complex that precedes formation of the cytotoxic *β*-pore employed by *Clostridium perfringens* to insult gastrointestinal epithelia in animals. The structures validate interactions between claudins and the cCpE domain alone found in previously determined structures but expand on these findings by illuminating the structure of nCpE during complex formation and the unique perturbations to claudin structure imposed by full CpE binding.^15,18-21^ In summary, we determined that hsCLDN-4 binds CpE variants with <10 nanomolar affinity and through EM showed that neither CpE nor claudin/CpE complexes form large assemblies *in vitro* (**Figure 1**). In detergents, small complexes do not assemble larger oligomers or pre-pores and is instead kinetically trapped. We show that X-ray crystallography, useful for obtaining structures of claudin/cCpE complexes, is severely limiting for small complex structure determination. The elongated structure of the small complex, natural hinge at the claudin/cCpE interface, and dynamics of detergent micelles explains this finding. Additionally, we show that these limitations can be overcome by cryo-EM with little modification to biochemical preparation. Cryo-EM reveals that hsCLDN-4 holds CpE in a single stable orientation at an ∼40° angle to the membrane plane (**Figures 2** and **3**). Both structures confirm that ECS2 and specifically the NPLVA^153^ motif within the ECH2-β5 loop drive high-affinity complex formation by penetrating a cleft in the cCpE domain of CpE (**Figure 4**).^15-17,21,31^ The two loops of ECS1 function in binding with gradually less importance where β1-2 binds the surface of the cCpE domain while cCpE’s 210-219 helix disrupts the β3-4 loop conformation. CpE-induced structural changes to hsCLDN-4 are thus relatively minor in ECS2 and become progressively more significant from β1-2 to β3-4 to β4-TM2 loops, which is altered the most (**Figure 4B**). Further, theoretical models shows that our experimentally determined small complexes can readily integrate into physiologically relevant pre-pores, which provides the potential biophysical procession from small complex to cytotoxic *β*-barrel pore (**Figure 5**). In full, this study provides evidence for how the gradient of claudin perturbation induced by CpE that culminates at the β4-TM2 loop of claudins could facilitate pre-cytotoxic pre-pore assembly.

How might CpE-induced structural alterations to claudins enable pre-pore assembly? It has been shown that ECH2 and the β4-TM2 loop, which may contain an ECH1 when unbound to toxin, direct cis interactions between claudins that influence tight junction morphology.^20,51-54^ It is thus curious that this ECH is unstructured in every claudin structure when bound to enterotoxin—a result of cCpE’s 210-219 helix disruption to ECS1.^15,18-21^ The unstructuring of ECH1 by CpE could thus lead to disabling of extracellular claudin cis interactions. Indeed, treatment of epithelial monolayers with cCpE has been shown to sequester claudin-4 from tight junctions to disrupt barrier function.^13^ Our study suggests another role of CpE alteration to claudin structure could be to promote the formation of unique extracellular claudin cis interactions (**Figure 5F**). We speculate that if claudins naturally form tight junction-favorable cis interactions using ECH2 and the β4-TM2 loop, that CpE binding could hijack these claudin interaction hubs and repurpose them to promote CpE oligomerization. In doing so, CpE binding would disfavor claudin integration into tight junctions and instead favor integration into CpE pores. The finding that cCpE removes claudin-4 from tight junctions could be a consequence of CpE-induced structural alterations to claudins that intend to drive CpE pore assembly, but cannot as nCpE was not present.^13^ If true, then CpEs binding pose may direct the directionality of pore assembly by keeping ECH2 rigid to form an interaction interface with the now structurally altered β4-TM2 loop to construct defined oligomeric pre-pores (**Figure S5**). In our proposed mechanism, claudins do not act solely as CpE receptors but instead as active participants in the CpE cytotoxic machinery. The choice of claudins as receptors by CpE would be a good one based on their propensity to assemble laterally in apical intestinal epithelium.

Based on this proposed mechanism, our structures and models imply that CpE uses cCpE to anchor to claudins but requires claudin/claudin interactions to drive CpE/CpE oligomerization and ultimately may use claudins to hold the complex together during the structural rearrangements nCpE undergoes during *β*-pore formation. Our kinetically trapped small complexes and inability to observe assemblies of CpE or small complexes in EM despite high concentrations, validates the idea that claudins actively participate in CpE oligomerization. We found previously that nCpE/nCpE interactions were not prevalent in the CpE_Tryp_ crystal structure.^29^ The cryo-EM structures reported here show that CpE is free to self-associate but does not—whereas claudins are barred from self-association by being encased in detergents. Without the possibility of claudin interactions, we observe that no larger complexes or pre-pores form despite CpEs availbility. To our knowledge, the idea that claudins actively enable CpE cytotoxicity has not been proposed before and thus requires experimental validation.

Our study also illuminates a potential role for trypsin in CpE cytotoxicity, which is known to activate toxicity three-fold but whose mechanism was unknown.^26-29^ We found earlier that trypsin removes the first 25 amino acids of CpE and that CpE_Tryp_ forms novel dimer interfaces *in crystallo* that are not observed in other CpE structures.^29^ The AF3 models of CpE pre-pores show that the short N-terminus of CpE_Tryp_ may interact with an adjacent cCpE domain to create of an interaction interface (**Figure 5E**). Because we show that N-term truncations of CpE bind claudin-4 with equivalent affinity, trypsin activation of CpE is not a result of changes in small complex formation but likely the subsequent formation of small complex oligomers (**Figure 1C**). The pre-pores architecture would place a full N-terminus between interaction interfaces (**Figures 5F** and **S5**). An entropically mobile and ∼3 kDa peptide could sterically shield CpE interactions required for pre-pore assembly. We thus hypothesize that removal of CpE’s N-terminus by trypsin eliminates this steric and entropic hindrance to facilitate small complex integration into functional pre-pores, and that this lower energy barrier to pre-pore assembly or the more favorable structural changes that result, activates CpE cytotoxicity.

What is the role of the membrane? Because the membranes properties alone or in conjunction with claudin interactions could facilitate CpE pore assembly, it is important to elucidate its role in CpE cytotoxicity. Our modeling provides potential insights. The AF3 hexamer, heptamer, octamer, and nonamer small complexes all show that individual claudins tilt at 30° angles within the membrane when forming conical dimer arrangements when analyzed using PPM 3.0 (**Figures 5** and **S5**).^55^ If true, these assemblies would positively curve the plasma membrane with the more small complexes comprising a pore, the greater the magnitude of curvature. In the pre-pore state, claudin-induced positive curvature would result in lipid packing defects focused at the center of the pore (**Figure S5**). CpE contains an amphipathic region between residues 81 and 108 and an amphipathic helix between residues 91 and 104 where Phe91, Phe95, Ile96, Val100, and Phe104 reside on one face of the helix and are shielded from solvent by nCpEs *β*-sheet (**Figure 4A**). The pattern of hydrophobic residues in the 81 and 108 region are characteristic of other *β*-pore forming toxins and it has been shown that mutants to the amphipathic helix reduce cytotoxicity via compromised membrane insertion.^56,57^ Synthesizing this information leads us to hypothesize that claudins, in addition to enabling pre-pore assembly through protein interactions, could bend membranes to recruit CpEs amphipathic domain, further governing pore assembly. Curved membranes are known to attract amphipathic helices and protein anchoring domains, and curvature could be the trigger that biophysically transitions pre-pores into *β*-pores.^58^ As before, small complexes in detergent would not be able to create an analogous environment, which may explain why we do not observe large assemblies, only kinetically trapped small complexes.

Synthesizing these and previous findings we propose the biophysical procession from small complex to cytotoxic *β*-barrel pore used by CpE during pathogenesis (**Figure 6**). In the small intestine, trypsin removes the N-terminus of CpE, which then uses its cCpE domain to target and bind claudin-4 selectively to form a small complex. CpE binding alters claudin-4 structure primarily at its β4-TM2 loop to prime it for novel cis interactions via extracellular and intracellular domains and dimerization. Claudin dimerization drives claudin oligomerization and in the process enables CpE/CpE interactions, one of which is formed by CpEs N-terminus as a specific result of trypsin cleavage, ultimately forming of a pre-pore complex. The conical arrangement and 30° tilt of claudins in a pre-pore resulting from the 40° angle that CpE is held at within each small complex, positively curves the plasma membrane. Membrane curvature creates lipid packing defects that recruit nCpEs amphipathic region to the membrane, triggering its structural reorganization and formation of a claudin-bound and membrane-permeable cytotoxic *β*-pore. The discrete steps in this process require experimental validation.

**Figure 6.**
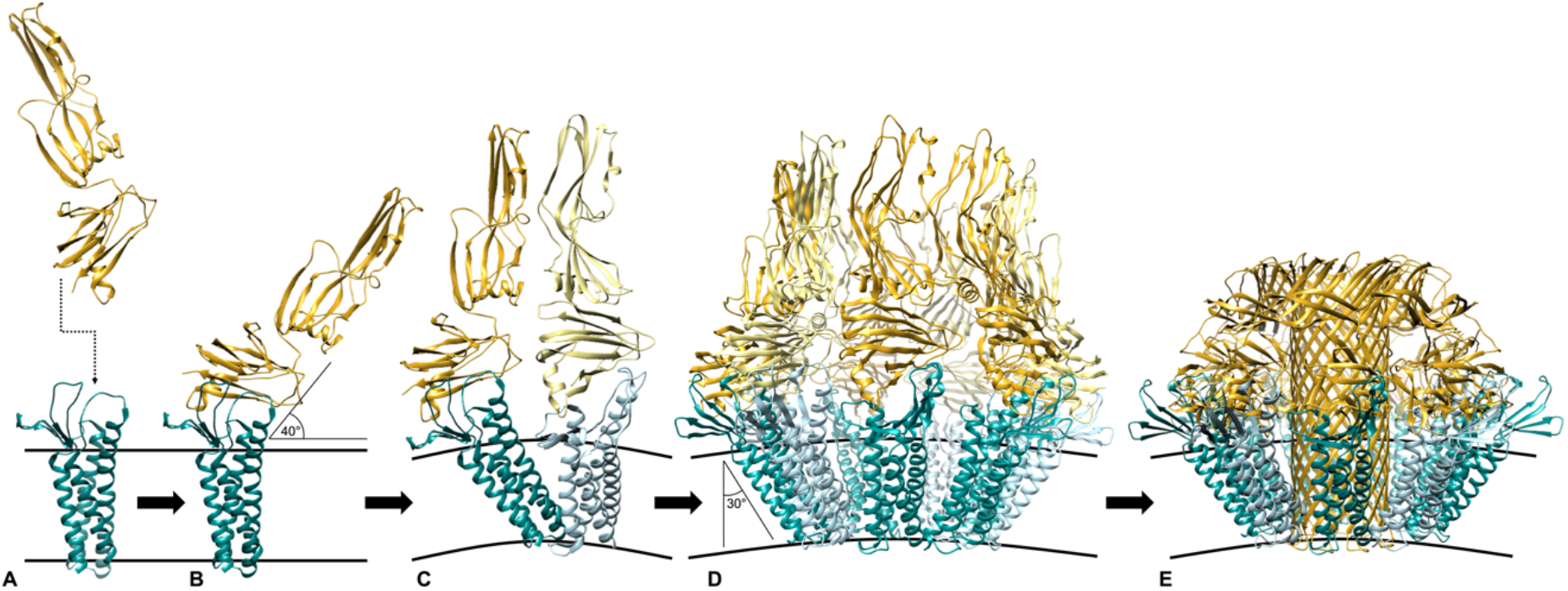
Biophysical Procession from Small Complex to Cytotoxic *β*-Barrel Pore. (A) CpE targets monomeric claudins out of tight junctions then form the (B) small complex. (C) Alteration of claudin ECS structure by CpE enables novel cis dimerization, which through claudin/claudin and CpE/CpE interactions, self-assemble into (D) an oligomeric pre-pore complex. The 40° angle CpE is held by claudins forces claudins to angle at 30° within membranes to prevent clashing, this causes membrane curvature. (E) Lipid packing defects are sensed by CpEs amphipathic domain, triggering reorganization of nCpE and membrane penetration into a cytotoxic *β*-pore similar to lysenin.^64^ Note: in D we used the AF3 nonamer model of the pre-pore to better align with nonameric lysenin.

Finally, this study provides a framework to obstruct small complex and pre-pore formation through structure-guided targeting of interaction interfaces with sFabs. First, we surmise it will be challenging to develop a sFab or other protein capable of binding the ECS of claudins or the cCpE of CpE with <1 nM to outcompete the 4 nM affinity that these proteins demonstrate for eachother.^15^ It could then be more tractable to target small complexes to prevent their integration into pre-pores. Using our models, we highlight the interfaces within oligomeric small complexes that could be bound by a sFab to potentially obstruct pre-pore assembly (**Figure 5G**). The targetable interfaces include all four in extracellular space, three formed by CpE and one by claudin. As 50 kDa proteins with six CDRs, sFabs are attractive molecules to target these interfaces as they could bind extensive and disparate regions of the small complex’s extracellular surface. A sFab that binds with high affinity at two or three interfaces simultaneously has potential as a therapeutic to kinetically trap the small complex per our modeling. If successfully, it may prevent its integration into pre-pores and/or the structural changes required for cytotoxic *β*-pore formation. Unexpectedly, COP-1 appears to obstruct CpE cytotoxicity to a degree. We predicted that because COP-1 does not interact with CpE, prevent small complex formation, and bind at any predicted pre-pore interface, that it would not inhibit CpE cytotoxicity (**Figure 5C**). However, our cell-based assay showed that 1 µM COP-1 decreased CpE cytotoxicity by an average of 9.8% compared to CpE alone (**Figure 5D**). This result demonstrates that either the AF3 models for pre-pore assembly are incorrect or that they are not and instead the structural changes from pre-pore to *β*-pore is hindered by the presence of COP-1. We found previously that COP-1 binds hsCLDN-4 small complexes in detergents with 15.5 nM affinity and that COP-1 can bind hsCLDN-4/cCpE complexes in lipid nanodiscs with ∼50 nM affinity.^45^ Yet we cannot verify whether COP-1 is bound to small complexes *in vivo* and thus the mechanism for COP-1s moderate inhibition of cytotoxicity requires further experiment. Ultimately, determining the structural bases of the CpE pre-pore and/or *β*-pore can rationally guide structure-based approaches aimed at inhibiting CpE cytotoxicity to treat *Clostridium perfringens* borne gastrointestinal diseases.

## Materials and Methods

### Protein expression and purification

All CpE variants (CpE, CpE NΔ33, and CpE_Tryp_) were expressed, purified, and produced as described previously.^15,29^ Briefly, all CpEs contained a C-terminal decahistidine tag preceded by a thrombin cleavage site (CpE-_His10_) and were expressed in insect cells. After cell lysis, and ultracentrifugation the supernatant was added to NiNTA resin (ThermoFisher), washed, and enterotoxins removed from resin via thrombin release for biochemical and structural studies. For biophysical studies, the histidine tag was retained by eluting CpE and CpE NΔ33 from NiNTA with 300 mM imidazole and removing excess imidazole by dialysis. CpE_Tryp_-_His10_ was produced by treating CpE-_His10_ with trypsin-immobilized resin (ProteoChem) using a 1:5 trypsin/CpE ratio (mass/mass) in 50 mM Tris pH 8.0, 200 mM NaCl, and 5% glycerol for 20 minutes at 4°C. Purity and homogeneity of CpE variants were assessed by SDS-PAGE and analytical SEC, and blue native-PAGE (Thermo-Fisher), using the manufacturers recommended protocols.

The hsCLDN-4 was expressed, purified, and produced as described previously.^45^ Briefly, hsCLDN-4-_His10_ was solubilized from membranes using n-dodecyl-β-D-maltopyranoside (DDM, Anatrace) and cholesteryl hemisuccinate Tris salt (CHS, Anatrace) to a final concentration of 1/0.1% (w/v), ultracentrifugated, then bound to NiNTA resin (ThermoFisher) and washed. For biochemical and biophysical applications, hsCLDN-4 was washed and released from the resin by thrombin. Purity and homogeneity of hsCLDN-4 was assessed by SDS-PAGE and analytical SEC using a Superdex 200 increase column equilibrated in SEC Buffer (20 mM Hepes pH 7.4, 100 mM NaCl, 1% glycerol, and 0.04% DDM). For crystallographic studies, hsCLDN-4 was solubilized from membranes using n-undecyl-β-D-maltopyranoside (UDM) CHS and washed on NiNTA in UDM with or without CHS. For cryo-EM studies, DDM solubilized hsCLDN-4 was washed with buffer containing 2,2-didecylpropane-1,3-bis-β-D-maltopyranoside (LMNG, Anatrace) and released by thrombin. Purity and homogeneity were assessed as before in their appropriate detergent systems.

COP-1 was expressed, purified, and produced as described previously.^45^ Briefly, COP-1 was expressed then extracted from bacterial cells in 0.01% DDM then purified using a protein L column (Cytiva). After affinity purification, COP-1 was dialyzed into SEC Buffer.

### Biophysical characterization

For biophysical analyses, 150 nM of CpE-_His10_ variants were immobilized on NiNTA (Dip and Read Sartorius) sensors and tested against 0-250 nM untagged hsCLDN-4 in BLI buffer (20 mM Tris pH 7.4, 100mM NaCl, 1% glycerol, and 0.03% DDM). BLI was performed at 25°C in 96-well black flat bottom plates (Greiner) using an acquisition rate of 5 Hz averaged by 20 using an Octet© R8 BLI System (FortéBio/Sartorius), with assays designed and setup using Blitz Pro 1.3 Software. Binding experiments consisted of sensor equilibration (30 seconds), loading (120 seconds), baseline (60 seconds), and association and dissociation (300 seconds each). Association and dissociation were fit to a 1:1 binding model using the Octet^®^ Analysis Studio (Sartorius). At the analyte concentrations used, no significant non-specific binding of hsCLDN-4 to NiNTA sensors were detected. Full results from a single experiment for each CpE variant and associated kinetic fits appear in **Table S1** and **Figure S1**.

### Assembly of small complexes

Post-NiNTA purified and tagless CpE and hsCLDN-4 were mixed using a 1.2 molar excess of CpE variant and analyzed by SEC using a Superdex 200 increase column equilibrated in SEC Buffer. Complexes that eluted prior to claudin alone were pooled and analyzed by SDS-PAGE and blue native-PAGE. Fractions containing both proteins were used for subsequent electron microscopy or crystallographic studies. For COP-1 small complexes, 1.3 molar excess COP-1 was added to hsCLDN-4 after CpE addition, then anti-sFab V_H_H nanobody (Nb) was added in 1.3 molar excess to COP-1.^46^ This complex was incubated for 20 minutes at room temperature then analyzed by SEC using a Superdex 200 increase column equilibrated in SEC Buffer. Complexes that eluted prior to small complexes were pooled and analyzed by SDS-PAGE. Fractions containing five proteins were used for subsequent electron microscopy or crystallographic studies.

### Crystallography of the small complex

Small complexes used for crystallization were assembled in ratios mentioned previously and eluted from SEC in Crystal Buffer (10 mM Hepes pH 7.4, 100 mM NaCl, 4% glycerol, and 0.08% UDM with or without 0.008% CHS), pooled, and concentrated to 15 mg/mL using a 50 kDa MWCO concentrator (Millipore). Using a Gryphon robot (Art Robbins Instruments) we added 80 µL of cocktail solution to the wells of 96-well 3-well Intelliplates (Art Robbins Instruments). 100 nL cocktail was added to drops then overlaid with 100-200 nL protein, sealed with crystal tape (Hampton), and incubated at 4, 10, or 15°C. Crystals were imaged using a CrysCam UV (Art Robbins Instruments) microscope. Crystals were harvested by hand removing them from 96- or 24-well plates using crystal mounts and loops (Mitegen). Some were vitrified after extraction from wells directly into well solutions supplemented with glycerol, ethylene glycol, or detergents; or streaking the crystals through 100% LV cryo oil (Mitegen); then were flash frozen in liquid nitrogen and stored until use. Diffraction was analyzed at Stanford Synchrotron Radiation Lightsource beamlines 12-1, 12-2, and 9-2; and Advanced Photon Source GM/CA beamlines 23ID-B and 23ID-D. Data was reduced using in-house beamline algorithms and programs or with XDS.^59^ Diffraction data from these experiments were visualized using adxv (Scripps: http://www.scripps.edu/tainer/arvai/adxv.html).

### Electron microscopy of small complexes

Details of negative stain EM imaging and data processing of small complexes conducted at the RTSF Cryo-EM Microscopy Facility at Michigan State University have been reported previously.^44^ For cryo-EM of the small complex, fractions containing both proteins from SEC in Cryo-EM Buffer (20 mM Hepes pH 7.4, 100 mM NaCl, and 0.003% LMNG) were pooled and concentrated to 4.4 and 8.0 mg/mL using a 50 kDa MWCO concentrator (Millipore). 3.5 μL of sample at each concentration were applied to UltrAuFoil 1.2/1.3 300 mesh (Quantifoil) grids that were glow-discharged for 60 seconds at 15 mA using a Pelco easiGlow (Ted Pella Inc) instrument. Grids were blotted for 3-5 seconds then plunge frozen into liquid ethane cooled by liquid nitrogen using an EM GP2 (Leica) plunge freezer at 4°C and 100% relative humidity. Grids were stored in liquid nitrogen then shipped overnight under cryogenic temperatures to the Pacific Northwest Cryo-EM Center (PNCC) for analysis. The grid that contained 8 mg/mL small complex blotted for 5 seconds was deemed superior and used for data collection. Cryo-EM data collection for the small complex was performed on a Titan Krios G3i (ThermoFisher) equipped with a Gatan K3 direct electron detector and BioContinuum HD GIF at PNCC. 7,100 movies were collected using SerialEM in counting mode at 129,000× magnification with a physical pixel size of 0.647 Å and super resolution pixel size of 0.3235 Å, defocus range of - 1.0 to -2.5 μm using a step of 0.1 μm, with a total dose of 60 electrons/Å^2^ fractionated over 59 total frames.

For cryo-EM of the COP-1 small complex, fractions containing five proteins from SEC in Cryo-EM Buffer were pooled and concentrated to 3.9 and 5.6 mg/mL using a 100 kDa MWCO concentrator (Millipore). Samples were shipped to the University of Chicago overnight on ice for vitrification and cryo-EM analysis. 3.5 μL of sample at each concentration were applied to UltrAuFoil 0.6/1.0 300 mesh (Quantifoil) grids that were glow-discharged for 30 seconds at 20 W using a Solarus 950 (Gatan) plasma cleaner. Grids were blotted for 3-5 seconds using a blot force of 2 then plunge frozen into liquid ethane cooled by liquid nitrogen using a Vitrobot Mark IV (ThermoFisher) plunge freezing apparatus at 8°C and 100% relative humidity. Grids were stored in liquid nitrogen before imaging then screened for thin ice and particle presence and distribution. The 5.6 mg/mL sample blotted for 5 seconds was used for data collection. Cryo-EM data collection for the COP-1 small complex was performed on a Titan Krios G3i (ThermoFisher) equipped with a Gatan K3 direct electron detector and BioQuantum GIF at the University of Chicago Advanced Electron Microscopy Core Facility (RRID:SCR_019198). 6,774 movies were collected using EPU (ThermoFisher) in CDS mode at 105,000× magnification with a super resolution pixel size of 0.827 Å and physical pixel size of 1.65 Å, defocus range of -0.9 to -2.1 μm using a step of 0.2 μm, with a total dose of 70 electron/Å^2^ fractionated over 50 total frames.

### Cryo-EM data processing and structure determination

Micrograph and particle processing for both complexes were performed in CryoSPARC.^60^ Patch-motion correction and patch-CTF correction were used to correct for beam-induced motion and calculate CTF parameters from the motion-corrected micrographs. Blob-based template picking was used first to generate templates that were subsequently used for template-based particle picking. Particles from template-based picking were subjected to multiple rounds of 2D classifications, followed by *ab initio* 3D reconstruction. For the small complex, heterogeneous refinement was followed by non-uniform refinement to yield the map with 4.0 Å global resolution. Of note, we found that default masking in non-uniform refinement after heterogeneous refinement made the maps visually worse and decreased resolution. We thus decreased the default dynamic mask start resolution to 5.5 Å after testing various parameters. Local refinement was run to try to improve map or resolution quality, but the resulting map was not altered or improved significantly from that of the non-uniform refined map (**Figure S2**). For the COP-1 small complex, heterogeneous refinement was followed by homogenous refinement of two classes, which were then 3D classified, re-extracted from micrographs, duplicates were removed and then subjected to several rounds of non-uniform refinement and 3D classification to yield the map with 2.83 Å global resolution. For both complexes, the final map local resolution was estimated by CryoSPARC’s algorithms using half maps. The summarized data processing workflows for both complexes are shown in **Figures S2** and **S3**, with statistics reported in **Table 1**.

For the small complex, the crystal structure of hsCLDN-4 bound to cCpE (PDB ID 7kp4) was placed into the cryo-EM density manually then fit into the map with Chimera.^47^ The crystal structure of CpE_Tryp_ (PDB ID 8u5f) was then superimposed onto cCpE and fit again into the map using Chimera, then rigid-body and real-space refined using Coot after replacing cCpE for CpE_Tryp_.^61^ For the COP-1 small complex, the cryo-EM structure of hsCLDN-4 bound to cCpE and COP-1/Nb (PDB ID 8u4v) was placed into the cryo-EM density and fit into the map with Chimera. Again, CpE_Tryp_ (PDB ID 8u5f) was superimposed on cCpE and fit into the map, then rigid-body and real-space refined using Coot after replacing cCpE for CpE_Tryp_. After model building both complexes in their respective maps using Coot, the final structural coordinates for each were generated after running phenix.real_space_refine against the final non-uniform refined CryoSPARC map in Phenix.^62^ The programs used to visualize and build the structures included Coot and Chimera, refinement in Phenix, and figures were made using Chimera— using the SBGrid Consortium Software Suite.^63^ **Table 1** shows refinement and validation statistics for both small complex structures.

### CpE *β*-Pore Modelling

We input the sequences of wild type hsCLDN-4 (residues 1-185, UniProt O14493) and CpE_Tryp_ (residues 27-319, UniProt P01558) into AlphaFold3 (AF3) using default parameters.^48^ The removal of hsCLDN-4s C-terminus was decided because AF cannot predict its structure with high accuracy as no experimental structure of it exists and we thought its inclusion would bias the predictions. We ran subsequent jobs searching for four to ten copies of each complex. After structural analysis of each output, tetramers, pentamers, and decamers were deems non-physiological due to their predicted assemblies. Structural coordinate outputs for claudin-bound hexamers, heptamers, octamers, and nonamers were downloaded and used for analysis.

### Cytotoxicity Assay

Methods were similar to those reported previously with slight modification.^15^ 2 mL of Sf9 cells (Expression Systems) were seeded on 6-well plates (Corning) at 80% confluency (1×10^6^ cells/mL) then hsCLDN-4_eGFP_ P1 baculovirus was added at a multiplicity of infection of 5 and left to incubate at 27°C. After 48 hours, 500 nM of CpE or CpE_Tryp_ (35 µg), or 1 µM (100 µg) COP-1 in BLI buffer were added to appropriate wells. For the COP-1 well, COP-1 was added first, incubated for 5 minutes, then CpE_Tryp_ was added. BLI buffer alone (vehicle) was added to wells not containing COP-1. The total volume of BLI buffer added to 2 mL media did not exceed 100 µL (5%). After 18 hours infection at 27°C, cells were imaged at 20x and 10x magnification using GFP filtered fluorescence microscope (Nikon). For cytotoxicity quantification, cells were then gently aspirated with a transfer pipette to remove adhering cells. 100 µL of cells were then added to 100 µL of 0.04% trypan blue and incubated for 5 minutes. After time, 10 µL of stained cells were transferred to counting chamber slides and counted immediately using a Countess 3 cell counter (ThermoFisher). Each well was counted in triplicate. Viability was determined by dividing the total number of live cells (unstained) by the total number of cells. Data with mean and standard error of the mean was plotted in GraphPad Prism 10 for macOS.

## Data availability

The cryo-EM structure of hsCLDN-4/CpE small complex has accession code XXXX in the Protein Data Bank (PDB) and cryo-EM maps of this complex have been deposited to the Electron Microscopy Data Bank (EMDB) under accession code EMD-XXXXX. The cryo-EM structure of the COP-1/Nb bound hsCLDN-4/CpE small complex has PDB accession code XXXX and EMDB accession code EMD-XXXXX.

## Acknowledgments

Research reported in this publication was supported by the National Institute of General Medical Sciences of the National Institutes of Health under Award Numbers R35GM138368 (to A.J.V.) and R01GM117372 (to A.A.K). The content is solely the responsibility of the authors and does not necessarily represent the official views of the National Institutes of Health. Grant support, cryo-EM training, and this material is based upon work supported by the National Science Foundation under Grant OIA-2131902 (to A.J.V.). We are grateful to the University of Chicago Advanced Electron Microscopy Core Facility (RRID:SCR_019198) for providing time and support for cryo-EM data collection. AJV is grateful to Craig Yoshioka, Claudia López, and Sean Mulligan at the Pacific Northwest Cryo-EM Center (PNCC) for training in cryo-EM acquired via NSF grant OIA-2131902. A portion of this research was supported by NIH grant U24GM129547 and performed at the PNCC at OHSU and accessed through EMSL (grid.436923.9), a DOE Office of Science User Facility sponsored by the Office of Biological and Environmental Research. AJV is also grateful to Benjamin Orlando at Michigan State University Department of Biochemistry and Molecular Biology for negative stain EM imaging and data processing and to the RTSF Cryo-EM Microscopy Facility at Michigan State University and Sundharraman Subramanian for providing time and support for negative stain EM data collection.

## Author contributions

A.J.V. designed the research; performed expression, purification, sample preparation for structure/function studies, and biochemical and biophysical characterization; crystallized small complexes and analyzed crystals via X-ray diffraction; processed cryo-EM data and determined cryo-EM structures; analyzed data; made figures; and wrote the paper. S.S.R. performed expression, purification, sample preparation for crystallography and was the first to crystallize the small complex. S.K.E. performed the sample vitrification, grid screening, and data collection of the COP-1 small complex. A.A.K. developed the sFab phage display library that discovered COP-1 and designed the sFab-related research.

