## Supplementary Information for "Cryo-EM Structures of *Clostridium perfringens* Enterotoxin Bound to its Human Receptor, Claudin-4"

ORCID: 0000-0001-8694-7681 (S.K.E.); 0000-0003-3174-9359 (A.A.K); 0000-0002-4222-7874 (A.J.V)

**This PDF file includes:**
Figures S1 to S6, Table S1

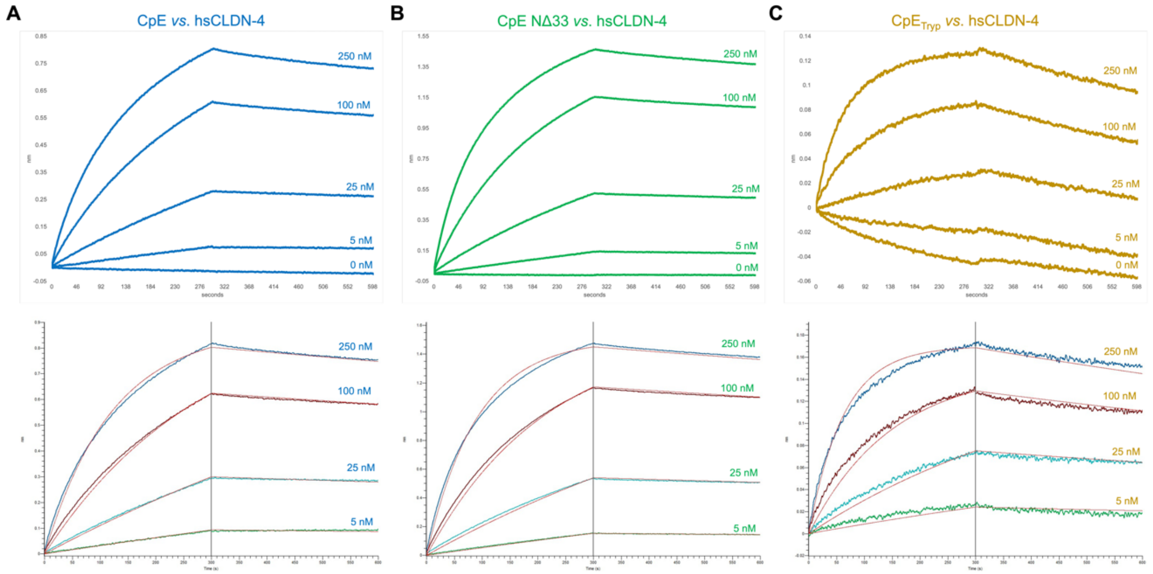


**Figure S1. BLI of CpE Variants *vs.* hsCLDN-4.** Binding of hsCLDN-4 analytes to (A) full-length CpE, (B) CpE NΔ33, and (C) CpE$_{Tryp}$. Top trace for each shows all five sensors from the experiment uncorrected as in **Figure 1C**. Bottom traces represent four ligand sensors subtracted from the 0 nM ligand sensor (control sample). Each ligand concentration is labeled and colored, with the kinetic fits depicted as red lines over the raw data. **Table S1** results were produced from the corrected data.

**Table S1. Binding of CpE Variants to hsCLDN-4.** BLI was used to measure binding kinetics and affinities of immobilized CpE variant ligands on NiNTA sensors *vs*. hsCLDN-4 analytes. Results represent a single experiment that was fit with a 1:1 binding model. The second-order association rate constant ($k_{on}$) and first-order dissociation rate constant ($k_{off}$) were used to calculate the equilibrium dissociation constant ($K_D$). The half-life ($t_{1/2}$) of a protein complex is also reported. **Figure S1** shows corrected BLI traces and associated fits that resulted in this data.

| Enterotoxin | $k_{on}$ (1/Ms) | $k_{off}$ (1/s) | $K_D$ (nM) | $t_{1/2}$ (min) |
|---|---|---|---|---|
| CpE | $3.2\times10^4$ | $2.4\times10^{-4}$ | 7.4 ± 0.1 | 48.9 |
| CpE NΔ33 | $4.4\times10^4$ | $2.1\times10^{-4}$ | 4.8 ± 0.1 | 55.3 |
| $CpE_{Tryp}$ | $6.2\times10^4$ | $5.0\times10^{-4}$ | 8.1 ± 0.1 | 23.0 |

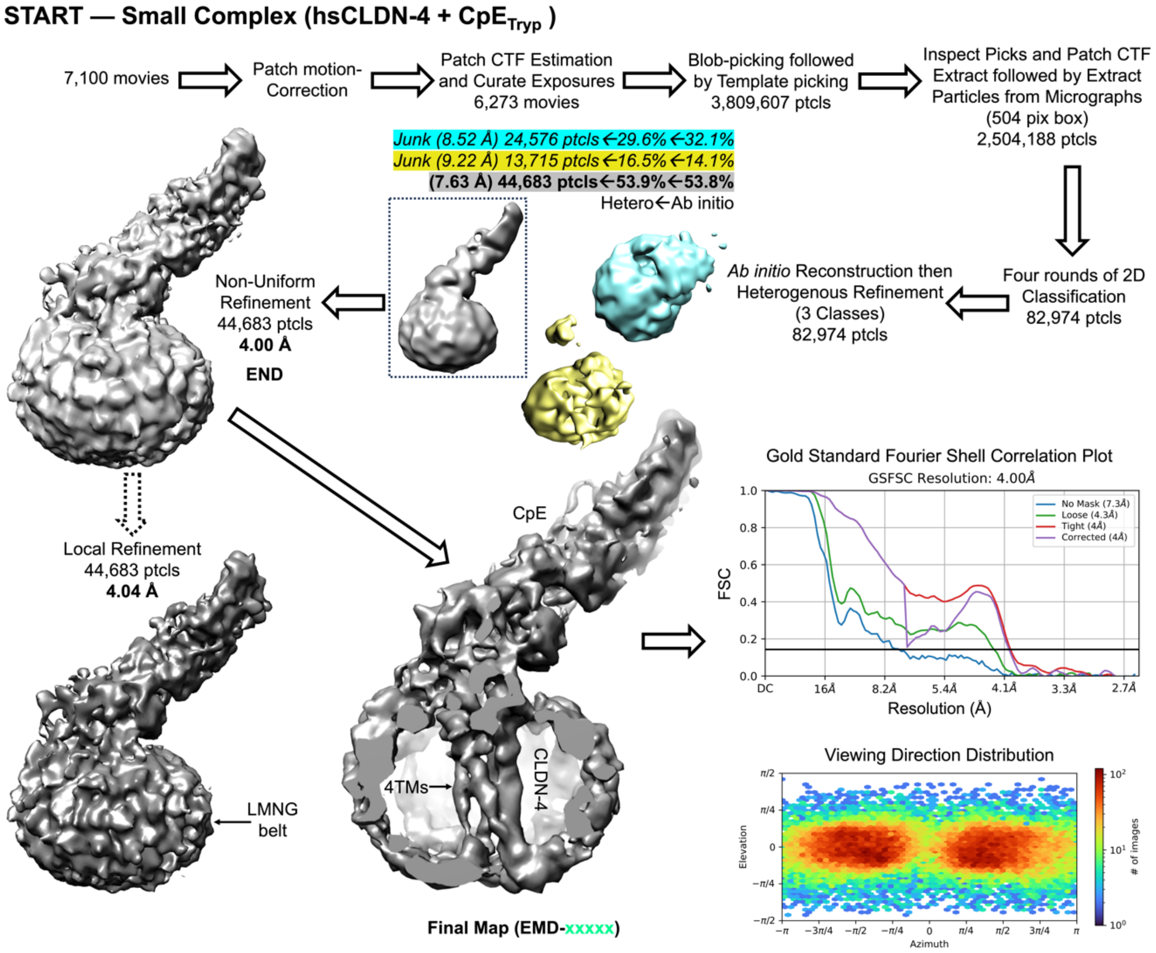


**Figure S2. Cryo-EM Data Processing Workflow for the Small Complex.** After four rounds of 2D classification, three *ab initio* reconstructions yielded one that resembled the small complex (gray). Non-uniform refinement and tuning of dynamic masking yielded the final 4.0 Å map. Attempts to improve the map quality by limiting contribution from the detergent belt using local resolution refinement did not improve the map significantly. Fourier Shell Correlation (FSC) curves from gold-standard refinement are shown with the 0.143 FSC cutoff indicated by line (black). Plots of the angular distribution of particles in the final refinement are shown below the FSC plot.

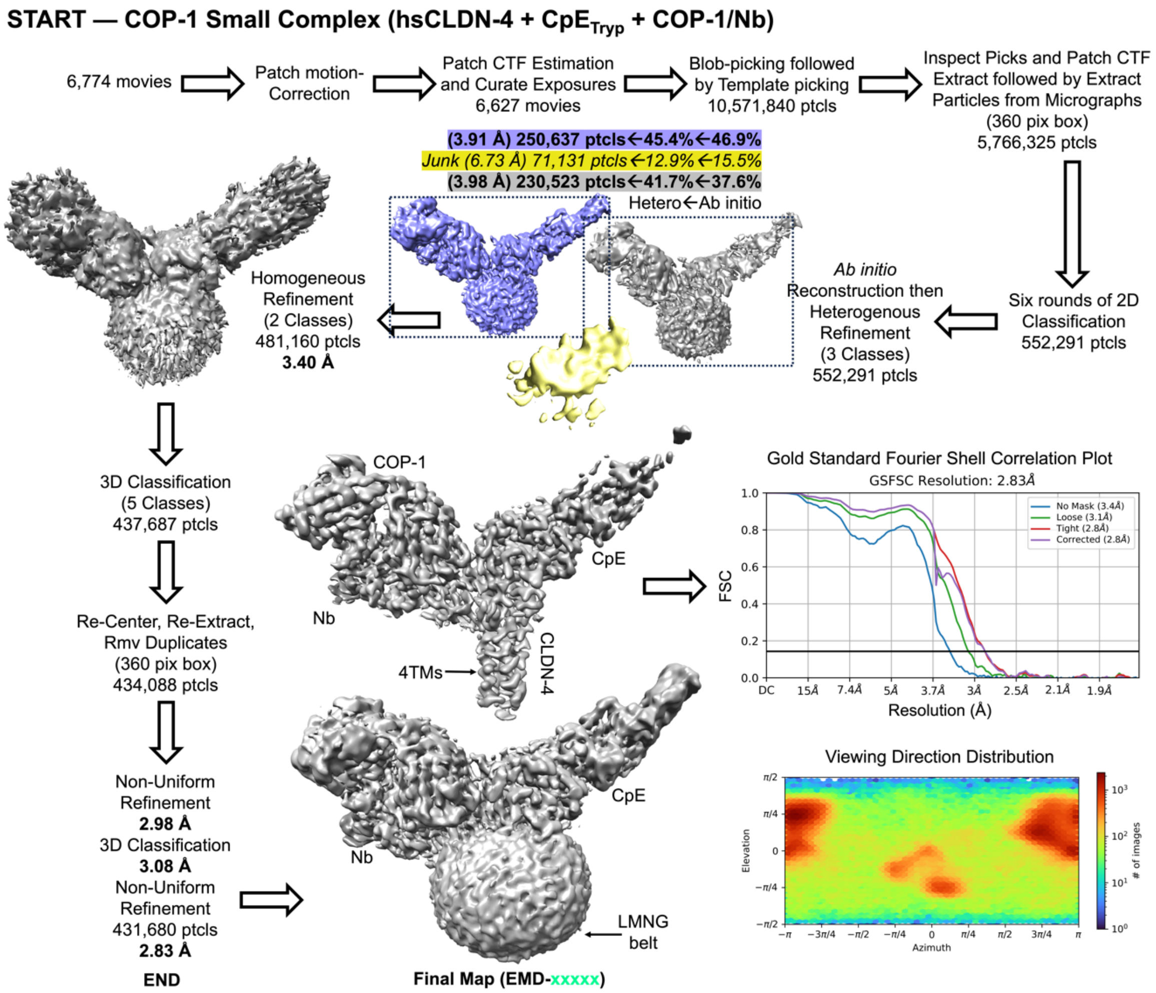


**Figure S3. Cryo-EM Data Processing Workflow for the COP-1 Small Complex.** After six rounds of 2D classification, three *ab initio* reconstructions yielded two that resembled the COP-1 small complex (purple and gray). Subsequent homogenous refinement, 3D classification, re-extraction and non-uniform refinement yielded the final 2.83 Å map. Tuning of the map contour level allows visualization of the claudin TM region contained within the detergent belt. Fourier Shell Correlation (FSC) curves from gold-standard refinement are shown with the 0.143 FSC cutoff indicated by line (black). Plots of the angular distribution of particles in the final refinement are shown below the FSC plot.

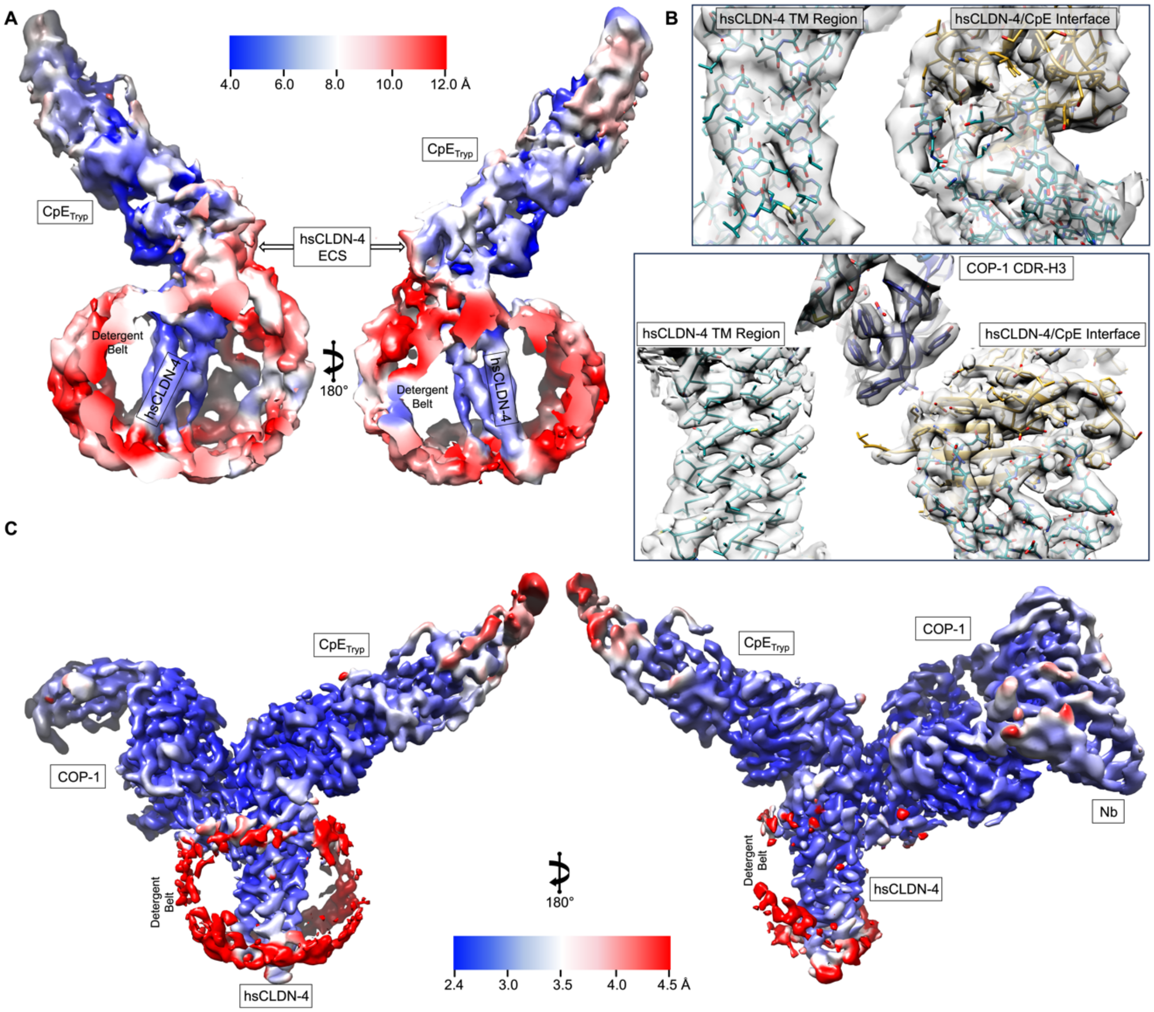


**Figure S4. Local Resolution Estimates and High-Resolution Features of Small Complexes.** (A) Local resolution estimates of the 4.0 Å small complex map. Regions of the map are colored from high (blue) and low (red) resolutions. (B) High resolution features of the maps for the small complex (top) and COP-1 small complex (bottom). hsCLDN-4 (teal), $CpE_{Tryp}$ (gold), and COP-1 (blue) are colored accordingly. Maps are translucent (grey) overlaid on the structures. (C) Local resolution estimates of the 2.8 Å COP-1 small complex map. Regions of the map are colored from high (blue) and low (red) resolutions.

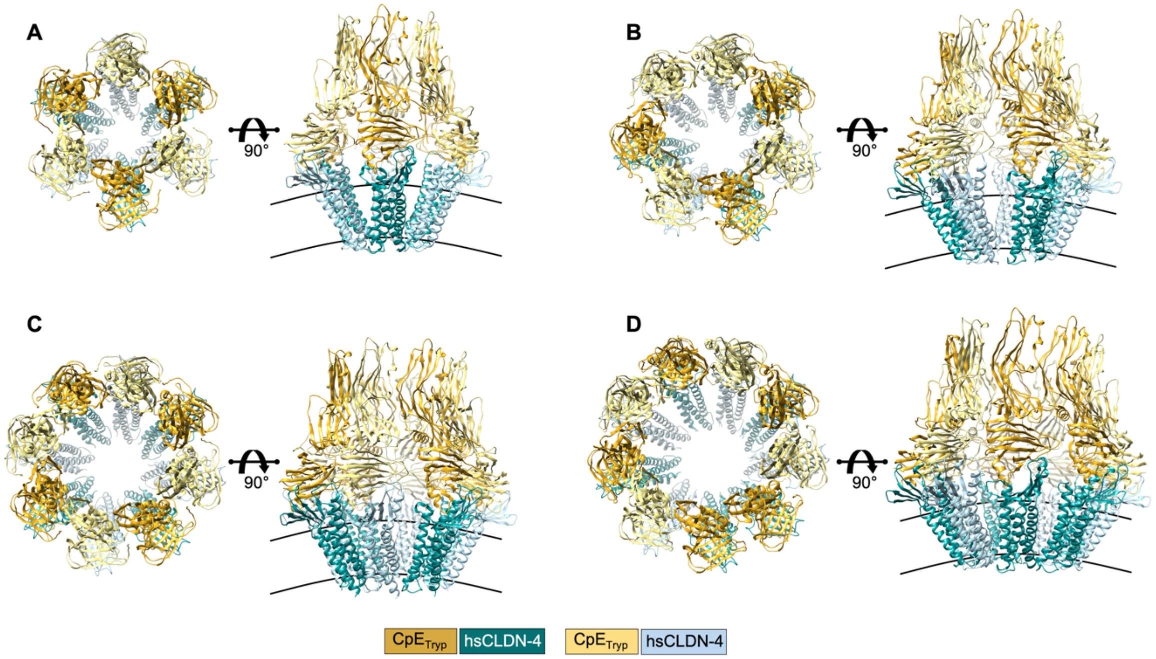


**Figure S5. AlphaFold 3 Predictions of Claudin-bound CpE Pre-Pore Oligomers.** We input the sequences for full-length hsCLDN-4 and $CpE_{Tryp}$ into the AF3 server searching for four to ten oligomeric complexes. AF3 predicted that (A) hexamers, (B) heptamers, (C) octamers, and (D) nonamers, could form oligomeric pre-pores organized in circular arrangements with a large central cavity. The organization and interfaces found in the hexamer were nearly identical to those from these other oligomers. Tetramers, pentamers, and decamers appeared non-physiological and were not investigated further.

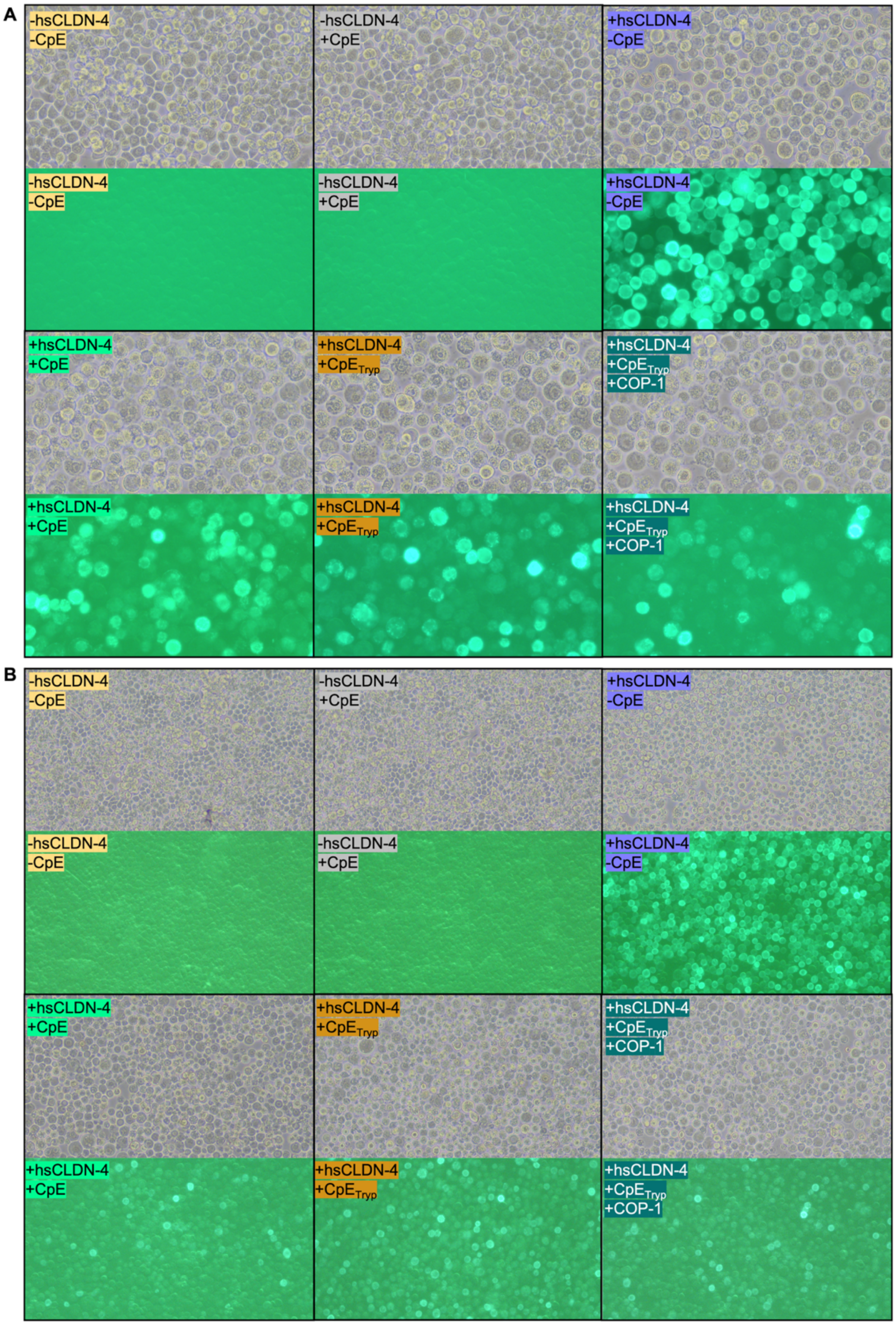


**Figure S6. CpE Cytotoxicity of Claudin-expressing Cells.** Sf9 insect cells expressing hsCLDN-4$_{eGFP}$ were treated with CpE, CpE$_{Tryp}$, or CpE$_{Tryp}$ and COP-1 then visualized using light and fluorescence microscopy at (A) 20x and (B) 10x magnification. Note that cells not expressing hsCLDN-4 exhibit no fluorescence and those expressing hsCLDN-4 and treated with CpE exhibit decreased fluorescence signals due to CpE-induced cell death. In the two control wells (-hsCLDN-4/-CpE and -hsCLDN-4/+CpE), undamaged cells appear mostly circular, of uniform size, and regularly shaped. In wells infected with hsCLDN-4$_{eGFP}$ baculovirus but not CpE (+hsCLDN-4/-CpE), cells appear larger than controls as a result of infection but still circular and regularly shaped with little morphological damage. In wells infected with hsCLDN-4$_{eGFP}$ baculovirus and CpE, morphological damage is shown by irregular shaped and much larger sized cells with decreased smoothness caused by membrane blebbing. Because this microscopy cannot quantify cell viability, these cells were stained, resulting in the data found in **Figure 5D**.
